# H3K4me2 orchestrates H2A.Z and Polycomb repressive marks in *Arabidopsis*

**DOI:** 10.1101/2025.05.26.656066

**Authors:** Takumi Noyori, Shusei Mori, Satoyo Oya, Haruki Nishio, Hiroshi Kudoh, Soichi Inagaki, Tetsuji Kakutani

**Author notes:** These authors contributed equally to this work (Takumi Noyori, Shusei Mori). Correspondence and requests for materials should be addressed to T.N., S.I, or T.K.

## Abstract

The dimethylation of histone H3 lysine 4 (H3K4me2) plays an important role in developmental phase transitions in plants, such as regeneration from the callus and the initiation of flowering. H3K4me2 in plants is correlated with transcriptionally repressed states, which can be accounted for by transcription-coupled active H3K4me2 demethylation, but the converse molecular pathway by which H3K4me2 represses transcription remains largely unexplored. Here, we show that H3K4me2 colocalizes with the facultatively repressive marks H2A.Z, H2A ubiquitination (H2Aub), and H3K27me3 in the gene body. Our genetic analyses reveal that H3K4me2 functions upstream but not downstream of these repressive marks. Interestingly, in genes with diurnal H3K4me2 oscillation, H3K4me2 oscillates in antiphase with transcription but in phase with H2A.Z. In addition, the genetic manipulation of H3K4me2 affects the oscillating profiles of H2A.Z, suggesting the efficient relay from H3K4me2 to H2A.Z. Notably, the diurnal oscillation of H2Aub is much weaker than that of H2A.Z despite the overall similarity in their distributions. These results suggest that H3K4me2 orchestrates H2A.Z and H2Aub with distinct dynamics. Our findings reveal that plants employ H3K4me2 to transmit transcriptional cessation into repressive chromatin states.

## Introduction

Methylation of histone H3 lysine 4 (H3K4) is a widely conserved modification among eukaryotes. H3K4 methylation has three states depending on the number of methyl groups: H3K4 monomethylation (H3K4me1), dimethylation (H3K4me2), and trimethylation (H3K4me3). H3K4 methylation generally marks actively transcribed genes; therefore, H3K4 methylation has been thought to play a role in activating transcription^1^. Indeed, H3K4me3 is strongly positively correlated with the transcription output of the genes in diverse species^2,3^. However, H3K4me2 is negatively correlated with transcription in plants, including the monocot rice and the eudicot *Arabidopsis thaliana*^2^. The removal of H3K4me2 by LYSINE-SPECIFIC DEMETHYLASE 1-LIKE 3 (LDL3) is necessary for the activation of genes involved in shoot regeneration from callus^4^. LDL3 binds to the phosphorylated C-terminal domain of RNA polymerase II and demethylates H3K4me2 in a transcription-coupled manner^5^. These findings suggest that transcription and H3K4me2 repress each other in plants^2–7^. While the effect of transcription on H3K4me2 is beginning to be understood mechanistically^5^, it is still unknown how H3K4me2 negatively regulates transcription.

Multiple histone methyltransferases in plants catalyze H3K4me1, H3K4me2, and H3K4me3 with distinct specificities^8,9^. In *A. thaliana*, ARABIDOPSIS TRITHORAX 3 (ATX3), ATX4, and ATX5 are mainly responsible for H3K4me2^8,9^. Although the loss of function of ATX3/4/5 or LDL3 alters H3K4me2 localization at thousands of genes, that does not result in widespread changes in gene expression^5,8^, suggesting that H3K4me2, as H3K4me1 and H3K4me3, functions depending on the developmental and chromatin contexts^9–11^.

The chromatin-based gene repression in the context of development has been well investigated in both plants and animals. The facultative heterochromatin is marked by the trimethylation of histone H3 lysine 27 (H3K27me3) and mono-ubiquitination of histone H2A (H2Aub), which are catalyzed by Polycomb repressive complex 2 (PRC2) and PRC1, respectively^12^. In *A. thaliana*, PRC1 acts upstream of PRC2, as the loss of PRC1 components affects the incorporation of H3K27me3, while PRC2 activity is generally not required for H2Aub^13^. A histone H2A variant, H2A.Z in the gene body region also has a repressive effect on transcription in cooperation with PRC^14–17^; H2A.Z can be ubiquitinated by PRC1^16^, and H2A.Z regulates H3K27me3 deposition in plants^18^. Interestingly, recent studies in *A. thaliana* demonstrated that the deposition of the repressive H2A.Z and H2Aub is promoted by H3K4me3, which is an active mark ^10,11^, and these contradictory trends are discussed in relation to bivalent chromatin modifications^10,11^. On the other hand, H3K4me2 is known to be negatively correlated with transcription^2,5^, but the possible connections between H3K4me2 and H2A.Z or H2Aub have not been tested.

To explore the molecular function of H3K4me2 in plants, we focused on the colocalization of H3K4me2 with H2A.Z, H2Aub, and H3K27me3. Using a genetic approach with mutants of genes encoding H3K4me2 methyltransferases and demethylases, we showed that an increase in H3K4me2 induces the accumulation of H2A.Z and H2Aub and that a decrease in H3K4me2 reduces H2A.Z, H2Aub, and H3K27me3. This pathway was one-way; H2A.Z, H2Aub, and H3K27me3 act downstream of H3K4me2. We also found that H3K4me2 diurnally oscillates in phase with H2A.Z but in antiphase with transcription, highlighting tight cooperation between H3K4me2 and H2A.Z in diurnal transcriptional regulation. In contrast, H2Aub does not oscillate diurnally, despite its similarity to H2A.Z in the static profiles and responses to H3K4me2. We propose that H3K4me2 coordinates H2A.Z and H2Aub on distinct timescales.

## Results

### H3K4me2 colocalizes with H2A.Z, H2Aub, and H3K27me3

To identify chromatin marks that are functionally linked to H3K4me2, we first explored chromatin modifications that colocalize with H3K4me2. Chromatin immunoprecipitation sequencing (ChIP-seq) in the wild type (WT) plants revealed that H2A.Z and H2Aub colocalized with H3K4me2 (Fig. 1a-e). These three chromatin marks were mainly accumulated in protein-coding genes and partially in non-protein-coding genes and pseudogenes, but did not accumulate in transposable element genes (Fig. 1a). Hereafter, we focused on chromatin marks in protein-coding genes. The distribution patterns of H3K4me2 within genes were similar to those of H2A.Z and H2Aub; these three marks mainly accumulated around the transcription start site (TSS), whereas some genes had broader accumulation of these marks throughout the gene bodies (Fig. 1a, d, e), which is consistent with previous reports on the localization patterns of H3K4me2^19^, H2A.Z^20^, and H2Aub^13,21^ in *A. thaliana*. H2A.Z and H2Aub levels were positively correlated with H3K4me2 levels across genes (Fig. 1b). When comparing accumulation levels among subregions within gene bodies, H2A.Z and H2Aub levels presented high correlations with H3K4me2 between the same subregion from the TSS to the transcription end site (TES) (Fig. 1c). These observations indicate that H2A.Z and H2Aub accumulate in the same genes and the same regions as H3K4me2. Consistent with previous reports^19,22^, H3K4me2 and H3K27me3 were colocalized only in a subset of genes (Fig. 1a, b, e). H3K4me2 levels in the genic region from the TSS to approximately 30% downstream of the TSS were negatively correlated with local H3K27me3 levels, whereas H3K4me2 levels further downstream were positively correlated with global H3K27me3 levels (Fig. 1c).

**Fig. 1.**
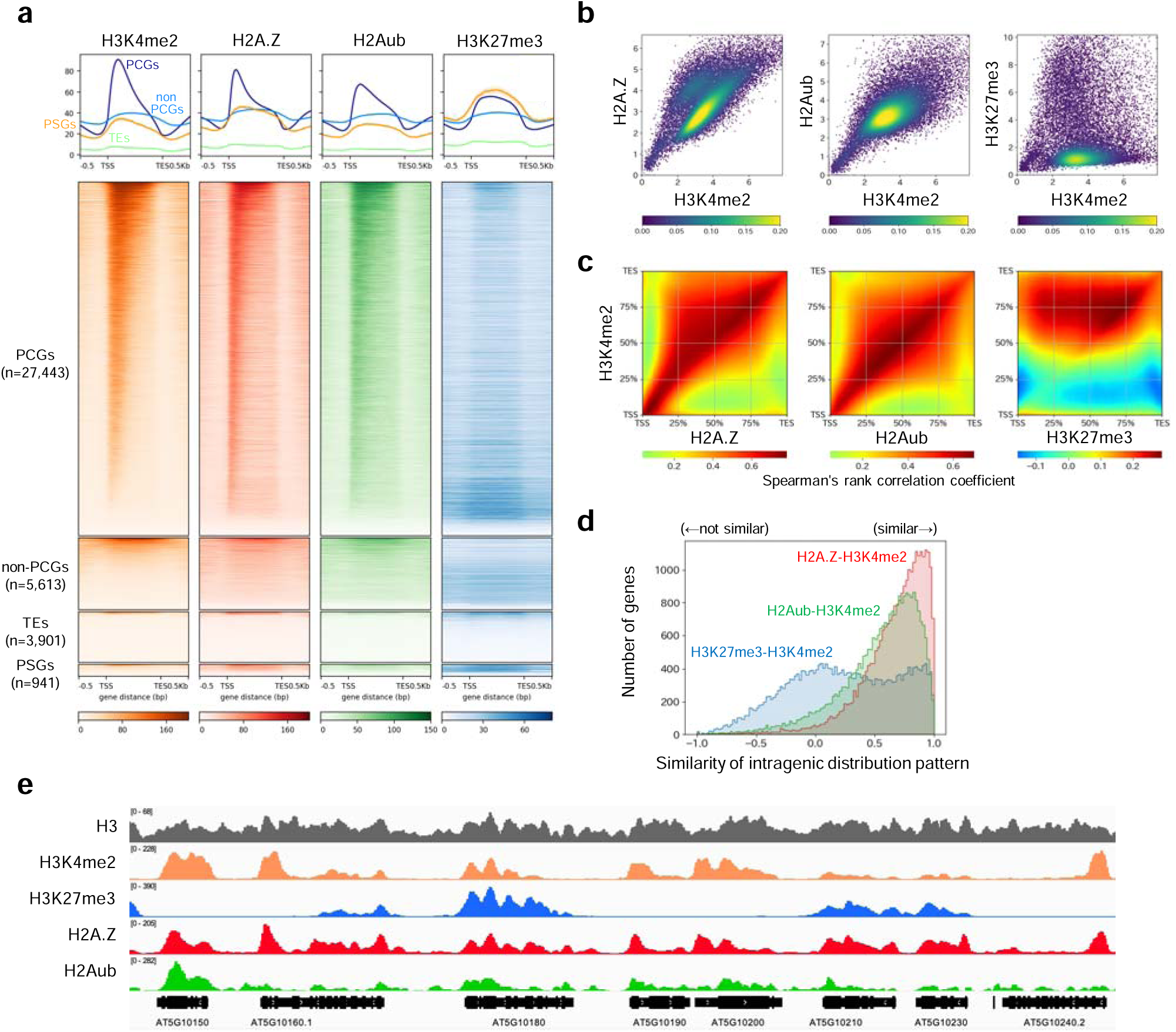
Genome-wide distribution of H3K4me2, H2A.Z, H2Aub, and H3K27me3 in wild type plants. **a**, Metaplots and heatmaps of H3K4me2, H2A.Z, H2Aub, and H3K27me3 in WT. All genes (n=37,898) are divided into 4 types: protein-coding genes (PCGs; n=27,443), non-protein-coding genes (non-PCGs; n=5,613), transposable element genes (TEs; n=3,901) and pseudogenes (PSGs; n=941). Genes are sorted in descending order by H3K4me2 levels. **b**, Scatter density plots showing the relationship between H3K4me2 levels and other chromatin mark levels (square root of RPKM). Each dot represents a protein-coding gene. Colors represent density. **c**, Heatmaps showing Spearman’s rank correlation coefficient between H3K4me2 and other chromatin marks (H2A.Z, H2Aub, and H3K27me3) calculated for each segmented gene region. Each gene region (TSS to TES) is divided into 100 equal parts. Higher coefficient values indicate greater similarity across all protein-coding genes between each pair of the 100 parts. **d**, Histograms showing the similarity of distribution pattern between H3K4me2 and H2A.Z (red), H3K4me2 and H2Aub (green), and H3K4me2 and H3K27me3 (blue) per gene. The similarity is calculated by Spearman’s rank correlation coefficient between read numbers mapped in 100 equally sized subregions of each gene. Higher coefficient values indicate similar intragenic distribution. **e**, A representative genome browser view of ChIP-seq signals of H3, H3K4me2, H3K27me3, H2A.Z, and H2Aub in WT.

Compared with H3K4me1 and H3K4me3, H3K4me2 showed greater positive correlations with H2A.Z and H2Aub (Supplementary Fig. 1a). H3K4me2 colocalized with H3K27me3 in many genes, whereas H3K4me3 was strongly exclusive to H3K27me3 (Supplementary Fig. 1a). Consistent with previous reports^13,16^, H2A.Z and H2Aub show strong colocalization, while their colocalization is modest for H3K27me3 (Supplementary Fig. 1b). Overall, we found strong colocalization of H3K4me2 with H2A.Z, H2Aub and, to a lesser extent, with H3K27me3.

### H2A.Z and H2Aub levels increase along with H3K4me2 in *ldl3*

The colocalization of H3K4me2 with H2A.Z, H2Aub, and H3K27me3 prompted us to investigate potential crosstalk among these chromatin marks. To determine whether H3K4me2 affects the distribution of H2A.Z, H2Aub, and H3K27me3, we perturbed the H3K4me2 pattern in plants using *ldl3* mutants. Previous studies have shown that a defect in the histone demethylase LDL3 causes an increase in H3K4me2 levels in thousands of actively transcribed genes^4,5^.

We performed ChIP-seq for H3K4me2, H2A.Z, H2Aub, and H3K27me3 in WT and *ldl3.* We defined genes with increased H3K4me2 in *ldl3* in both independent experiments as LDL3-target genes (n=7,115; Fig. 2a). Focusing on LDL3-target genes, H2A.Z and H2Aub levels increased in *ldl3*, especially in the gene body, as did H3K4me2 (Fig. 2b-d). The increase in H2A.Z and H2Aub levels in *ldl3* were proportional to that in H3K4me2 levels (Fig. 2e). To examine whether the change in H2A.Z and H2Aub levels in *ldl3* is due to changes in the transcription of their regulators, the expression of H2A.Z- and H2Aub-related genes in *ldl3* were compared with those in WT. A reanalysis of our previous mRNA-seq data^5^ revealed that the expression of H2A.Z-related genes (e.g., *HTA8*) and H2Aub-related genes (e.g., *AtBMI1A*) did not significantly change in *ldl3* (Supplementary Fig. 2d). These results support the idea that the increase in H3K4me2 itself, rather than secondary effects of the *ldl3* mutation, induces the accumulation of H2A.Z and H2Aub. H3K27me3 slightly increased in *ldl3* in LDL3-target genes, although H3K27me3 levels were generally low in genes with highly increased H3K4me2 in *ldl3* (Fig. 2b-e).

**Fig. 2.**
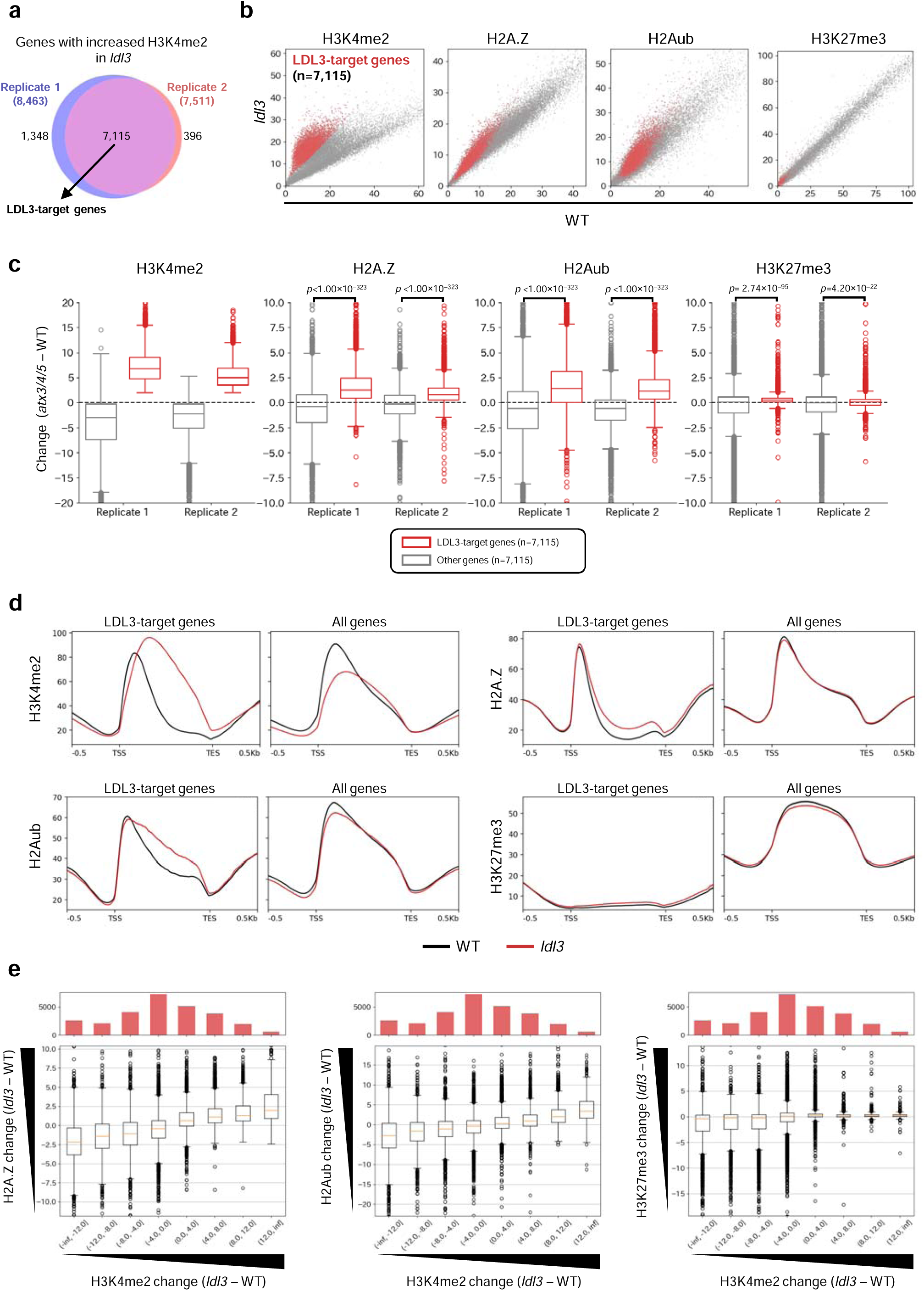
H2A.Z and H2Aub increase along with H3K4me2 in *ldl3*. **a-e,** ChIP-seq for H3K4me2, H2A.Z, H2Aub, and H3K27me3 in WT and *ldl3*. **a**, Overlap of genes with increased H3K4me2 in *ldl3* compared to WT (H3K4me2 change (*ldl3*–WT) > 2) between two biological replicates. The overlapping genes are defined as “LDL3-target genes (n=7,115)”. **b**, Scatter plots showing the levels of H3K4me2, H2A.Z, H2Aub, and H3K27me3 comparing WT and *ldl3*. Each dot represents a protein-coding gene. Red dots indicate LDL3-target genes (n=7,115). **c**, Box plots showing the change in H3K4me2, H2A.Z, H2Aub, and H3K27me3 (*ldl3*–WT) in LDL3-target genes (n=7,115) and other genes (7,115 genes were randomly selected from genes other than LDL3-target genes). The p-values from two-sided Mann-Whitney U tests are shown. **d,** Metaplots showing the averaged distribution of H3K4me2, H2A.Z, H2Aub, and H3K27me3 in WT (black) and *ldl3* (red) in LDL3-target genes (n=7,115) and all protein-coding genes (n=27,443). **e,** Box plots showing the relationship between the change in H3K4me2 and the change in H2A.Z, H2Aub, and H3K27me3 in *ldl3* compared to WT. Histograms above the boxplots indicate the number of genes in each category of H3K4me2 change. The values of scatter plots, metaplots, and box plots represent RPKM. An additional series of the ChIP-seq analyses for **b**, **d**, and **e** showed reproducible results (Supplementary Fig 2a-c).

### H2A.Z, H2Aub, and H3K27me3 levels decrease along with H3K4me2 in *atx3/4/5*

As a complementary approach, we next examined whether a decrease in H3K4me2 levels affects the localization of H2A.Z, H2Aub, and H3K27me3 using triple mutants of the H3K4 methyltransferases ATX3, ATX4, and ATX5. H3K4me2 drastically decreased in *atx3/4/5*, which is consistent with our previous study^9^. The 5,842 genes with decreased H3K4me2 levels in *atx3/4/5* in both independent experiments were defined as ATX3/4/5-target genes (Fig. 3a). Notably, in ATX3/4/5-target genes, H2A.Z, H2Aub, and H3K27me3 levels also decreased in *atx3/4/5* (Fig. 3b-d). The decrease in H2A.Z, H2Aub, and H3K27me3 levels in *atx3/4/5* were proportional to that in H3K4me2 levels (Fig. 3e). We confirmed that the expression of H3K27me3-related genes (e.g., *CURLY LEAF* (*CLF*)), H2A.Z-related genes, and H2Aub-related genes did not significantly change in *atx3/4/5* (Supplementary Fig. 3d). Taken together, these results demonstrated that H3K4me2 positively regulates the accumulation of H2A.Z, H2Aub, and H3K27me3 in the same region.

**Fig. 3.**
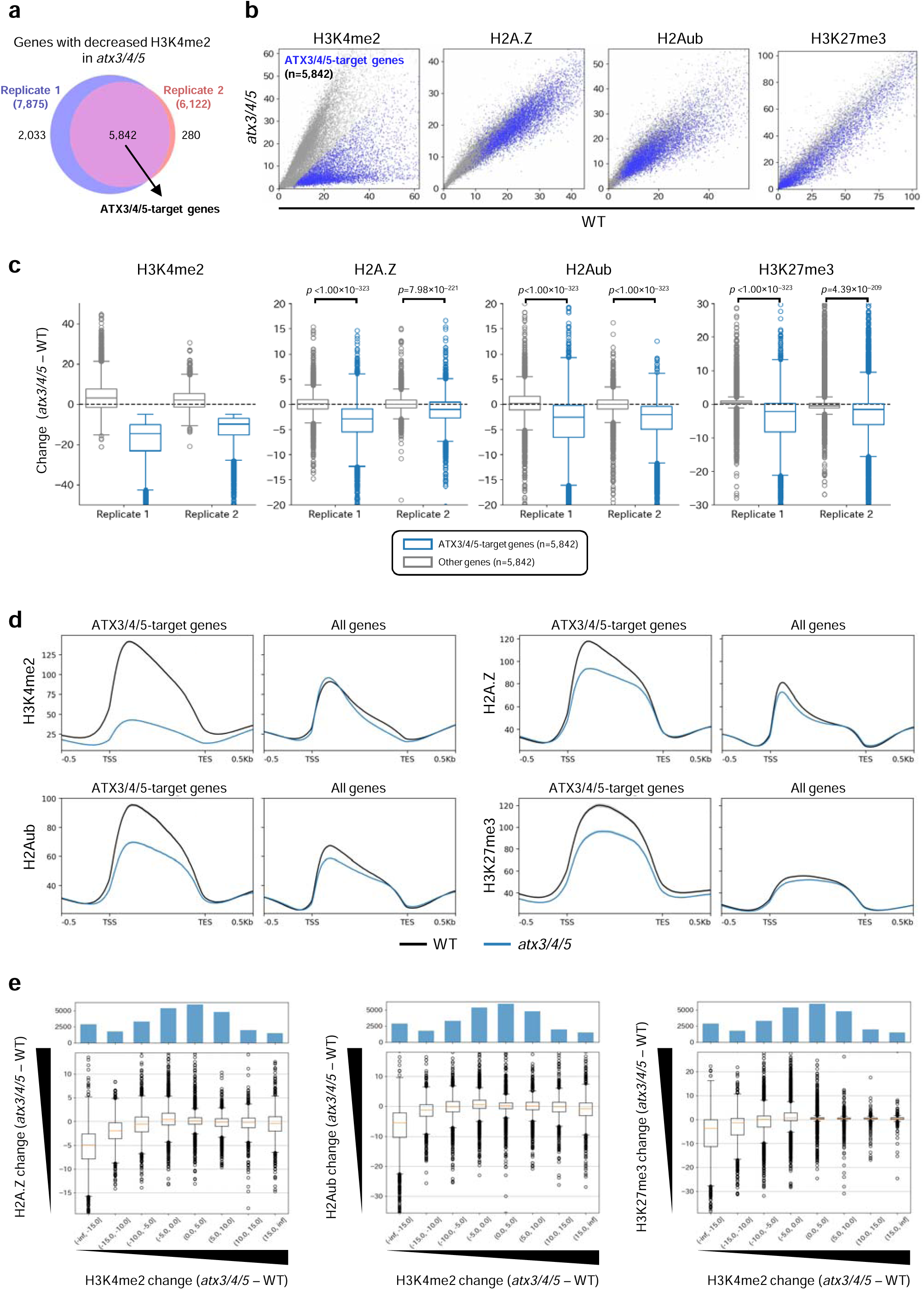
H2A.Z, H2Aub, and H3K27me3 decrease along with H3K4me2 in *atx3/4/5*. **a-e,** ChIP-seq for H3K4me2, H2A.Z, H2Aub, and H3K27me3 in WT and *atx3/4/5*. **a**, Overlap of genes with decreased H3K4me2 in *atx3/4/5* compared to WT (H3K4me2 change (*atx3/4/5*–WT) < –5) between two biological replicates. The overlapping genes are defined as “ATX3/4/5-target genes (n=5,842)”. **b**, Scatter plots showing the levels of H3K4me2, H2A.Z, H2Aub, and H3K27me3 comparing WT and *atx3/4/5*. Each dot represents a protein-coding gene. Blue dots indicate ATX3/4/5-target genes (n=5,842). **c**, Box plots showing the change in H3K4me2, H2A.Z, H2Aub, and H3K27me3 (*atx3/4/5*–WT) in ATX3/4/5-target genes (n=5,842) and other genes (5,842 genes were randomly selected from genes other than ATX3/4/5-target genes). The p-values from two-sided Mann-Whitney U tests are shown. **d,** Metaplots show the averaged distribution of H3K4me2, H2A.Z, H2Aub, and H3K27me3 in WT (black) and *atx3/4/5* (blue) in ATX3/4/5-target genes (n=5,842) and all protein-coding genes (n=27,443). **e,** Box plots showing the relationship between the change in H3K4me2 and the change in H2A.Z, H2Aub, and H3K27me3 in *atx3/4/5* compared to WT. Histograms above the boxplots indicate the number of genes in each category of H3K4me2 change. The values of scatter plots, metaplots, and box plots represent RPKM. An additional series of the ChIP-seq analyses for **b**, **d**, and **e** showed reproducible results (Supplementary Fig. 3a-c).

### Alterations in H3K27me3, H2A.Z, or H2Aub levels do not affect H3K4me2

We then tested the converse effect. First, a quintuple mutant of genes encoding Jumonji-C-domain-containing proteins (JMJs) with H3K27me3 demethylase activity, i.e., JMJ30, JMJ32, JMJ11/EARLY FLOWERING 6 (ELF6), JMJ12/RELATIVE OF EARLY FLOWERING (REF6), and JMJ13^23,24^, and a single mutant of the gene encoding the H3K27me3 methyltransferase CLF^25^, were used to investigate whether H3K27me3 regulates H3K4me2. The *jmj30 jmj32 elf6 ref6 jmj13* quintuple mutant exhibited ectopic accumulation of H3K27me3, but H3K4me2 did not change (Fig. 4a, b; Supplementary Fig. 4a). In addition, the decrease in H3K27me3 levels in thousands of genes in *clf* did not cause any change in H3K4me2 patterns (Fig. 4c, d; Supplementary Fig. 4b). These results demonstrate that altering H3K27me3 levels does not affect the localization patterns of H3K4me2.

**Fig. 4.**
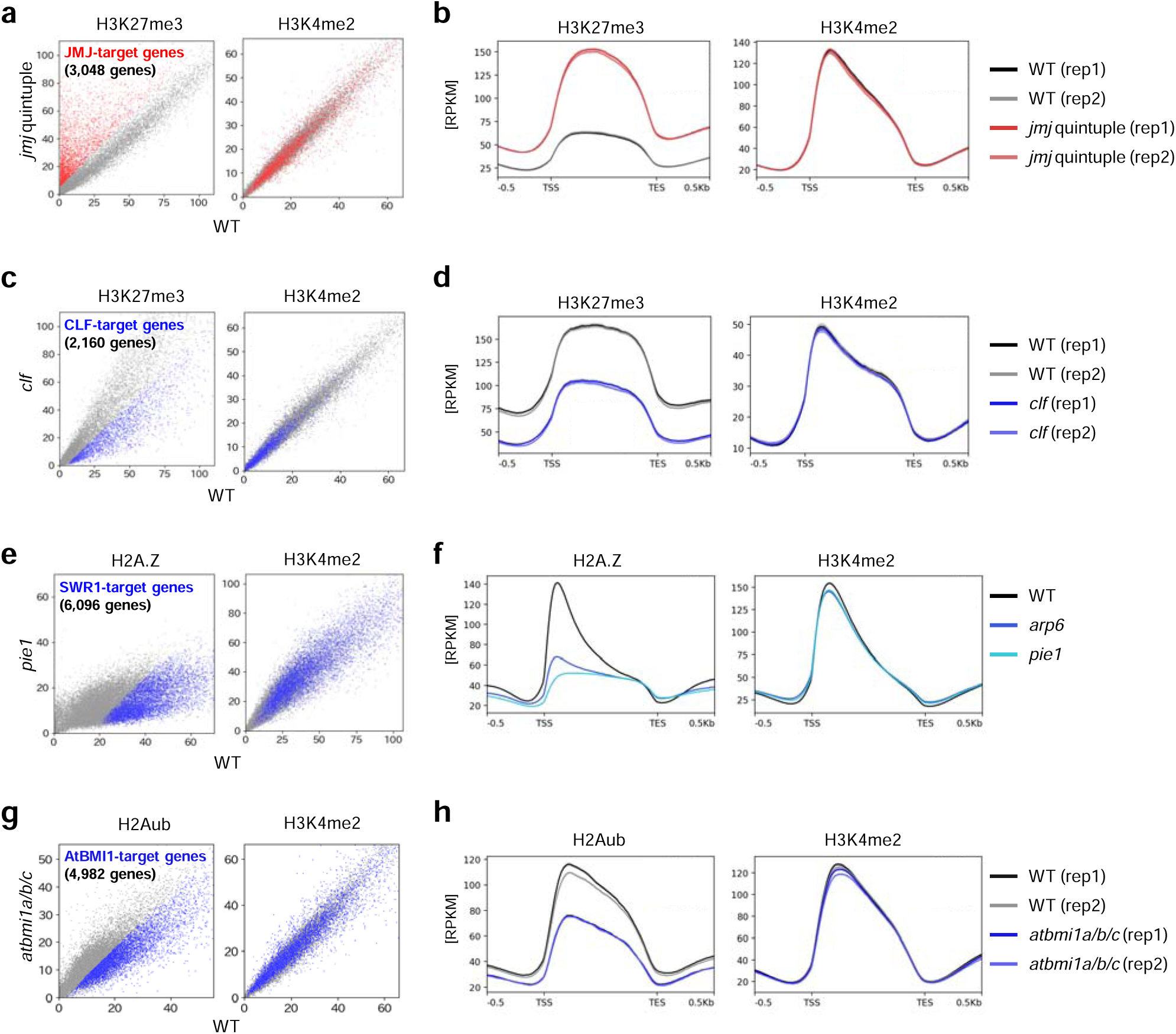
Alteration in H3K27me3, H2A.Z, or H2Aub levels do not affect H3K4me2. **a**, Scatter plots showing H3K27me3 and H3K4me2 levels comparing *jmj30 jmj32 elf6 ref6 jmj13* quintuple mutants (*jmj* quintuple) and WT. Each dot represents a protein-coding gene. Red dots indicate JMJ-target genes (n=3,048, as shown in Supplementary Fig. 4a). **b**, Metaplots showing the averaged distribution of H3K27me3 and H3K4me2 in JMJ-target genes (n=3,048) in WT and *jmj* quintuple. **c**, Scatter plots showing H3K27me3 and H3K4me2 levels comparing *clf* and WT. Each dot represents a protein-coding gene. Blue dots indicate CLF-target genes (n=2,160, as shown in Supplementary Fig. 4b). **d**, Metaplots showing the averaged distribution of H3K27me3 and H3K4me2 in CLF-target genes (n=2,160) in WT and *clf*. **e**, Scatter plots showing H2A.Z and H3K4me2 levels comparing *pie1* and WT. Each dot represents a protein-coding gene. Blue dots indicate SWR1-C-target genes (n=6,096, as shown in Supplementary Fig. 5c). The regions from TSS to TSS+500bp were analyzed. **f**, Metaplots showing the averaged distribution of H2A.Z and H3K4me2 in SWR1-C-target genes (n=6,096) in WT, *arp6*, and *pie1*. **g**, Scatter plots showing H2Aub and H3K4me2 levels comparing WT and *atbmi1a/b/c*. Each dot represents a protein-coding gene. Blue dots indicate AtBMI1-target genes (n=4,982, as shown in Supplementary Fig. 6a). **h**, Metaplots showing the averaged distribution of H2Aub and H3K4me2 in AtBMI1-target genes (n=4,982) in WT and *atbmi1a/b/c*. The values of scatter plots and metaplots represent RPKM.

To test the effect of H2A.Z to H3K4me2, we used the mutants of *PHOTOPERIOD-INDEPENDENT EARLY FLOWERING 1* (*PIE1*) and *ACTIN-RELATED PROTEIN 6* (*ARP6*), which encode the components of SWI2/SNF2-Related 1 chromatin remodeling complex (SWR1-C) responsible for H2A.Z deposition^26^. As most H2A.Z peaks were in the TSS to TSS+500 bp region of genes (Supplementary Fig. 5a), H2A.Z and H3K4me2 in this region were investigated. H2A.Z changes in *arp6* and *pie1* were highly correlated (Spearman’s rank correlation coefficient ρ=0.888; Supplementary Fig. 5b); therefore, we defined SWR1-C-target genes (n=6,096) as the overlap of genes with decreased H2A.Z levels in both *arp6* and *pie1* (Supplementary Fig. 5c). Although H3K4me2 levels were slightly disturbed in SWR1-C-target genes in *pie1* and *arp6*, the overall pattern of H3K4me2 was unchanged, and there was no clear correlation between H2A.Z changes and H3K4me2 changes in both *arp6* and *pie1* (Fig. 4e, f; Supplementary Fig. 5d, e).

To address whether a decrease in H2Aub levels affects H3K4me2 levels, we analyzed the effects of the loss of AtBMI1A, AtBMI1B, and AtBMI1C, which are components of the PRC1 complex responsible for the monoubiquitination of H2A^27–29^. We performed ChIP-seq for H3K4me2 and H2Aub in the *atbmi1a atbmi1b atbmi1c* triple mutant (*atbmi1a/b/c*). Despite the changes in H2Aub levels in AtBMI1-target genes, H3K4me2 levels did not change in *atbmi1a/b/c* (Fig. 4g, h; Supplementary Fig. 6a).

We also investigated the interplay between H2A.Z and Polycomb repressive marks. A previous study revealed that H3K27me3 decreases in *pie1*^18^. We found that H2A.Z did not change in the *jmj* quintuple mutant, *clf*, or *atbmi1a/b/c*, indicating that Polycomb repressive marks do not affect the deposition of H2A.Z (Supplementary Fig. 4c-f, Supplementary Fig. 6b-c).

Overall, we found that H3K4me2 colocalized with three transcriptionally repressive chromatin marks, H2A.Z, H2Aub, and H3K27me3. These chromatin marks responded proportionally to the changes in H3K4me2 (Fig. 2, Fig. 3) but not vice versa (Fig. 4), demonstrating that they act downstream of H3K4me2.

### H2A.Z oscillates in phase with H3K4me2 in the context of the circadian oscillation

We showed that H3K4me2 regulates the accumulation of H2A.Z, H2Aub, and H3K27me3 on chromatin. However, changes in H2A.Z and H2Aub levels in *ldl3* and *atx3/4/5* were lower than those in H3K4me2 (Fig. 2, Fig. 3), whereas H2A.Z and H2Aub strongly colocalized with H3K4me2 in WT (Fig. 1). This implies that there are unknown common factors upstream of H3K4me2, H2A.Z, and H2Aub, in addition to the regulation of H2A.Z and H2Aub by H3K4me2. Furthermore, the timescale over which H2A.Z, H2Aub, and H3K27me3 change after changes in H3K4me2 is still unknown. To address these issues, we analyzed the circadian oscillation of these chromatin marks in the WT plants. In *A. thaliana*, H3K4me2 levels were reported to show circadian oscillation in core clock genes such as *LATE ELONGATED HYPOCOTYL* (*LHY*), *CIRCADIAN CLOCK ASSOCIATED 1* (*CCA1*), and *TIMING OF CAB EXPRESSION 1* (*TOC1*)^6^. At least in the core clock genes, H3K4me2 diurnally oscillates in antiphase with transcription^6^. This let us consider the possibility that H2A.Z, H2Aub, and H3K27me3 also oscillate following H3K4me2 oscillation and that these chromatin marks synergistically regulate transcriptional oscillation. Because it has already been reported that H3K27me3 does not diurnally oscillate in *Arabidopsis* species^6,30^, we focused on H3K4me2, H2A.Z, and H2Aub and investigated the genome-wide circadian oscillation of these chromatin marks.

ChIP-seq for H3, H3K4me2, H3K4me3, H2A.Z, and H2Aub and mRNA-seq were performed using WT seedlings sampled at Zeitgeber Time 0 (ZT0), ZT6, ZT12, and ZT18 after growth for 14 days under a 12-h light/12-h dark cycle (Fig. 5a). We identified 781 genes with H3K4me2 oscillation (called “H3K4me2 circadian oscillation genes (COGs)”), 1,180 genes with H3K4me3 oscillation (“H3K4me3 COGs”), 345 genes with H2A.Z oscillation (“H2A.Z COGs”) and 29 genes with H2Aub oscillation (“H2Aub COGs”) (Fig. 5b). H3K4me2 COGs significantly overlapped with H2A.Z COGs, as well as H3K4me3 COGs and mRNA COGs (Fig. 5c). To dissect the temporal relationship between H3K4me2 and other chromatin marks, H3K4me2 COGs were classified into four gene groups on the basis of the time when H3K4me2 had reached its maximum value: genes with ZT0-peak (n=337), ZT6-peak (n=149), ZT12-peak (n=209), and ZT18-peak (n=86). For all ZT-peak groups, H2A.Z oscillated in phase with H3K4me2, whereas H3K4me3 and mRNA oscillated in antiphase with H3K4me2 (Fig. 5d). Consistent with this observation, the temporal change in H3K4me2 was positively correlated with that in H2A.Z and negatively correlated with that in H3K4me3 and mRNA for most of the H3K4me2 COGs, including core clock genes (Fig. 5e; Supplementary Fig. 7). On the other hand, H2Aub oscillation was much weaker and not synchronous with H3K4me2 (Fig. 5d, e; Supplementary Fig. 7). These results uncovered the difference between H2A.Z and H2Aub in the diurnal oscillation, in addition to the tight temporal association of H2A.Z with H3K4me2.

**Fig. 5.**
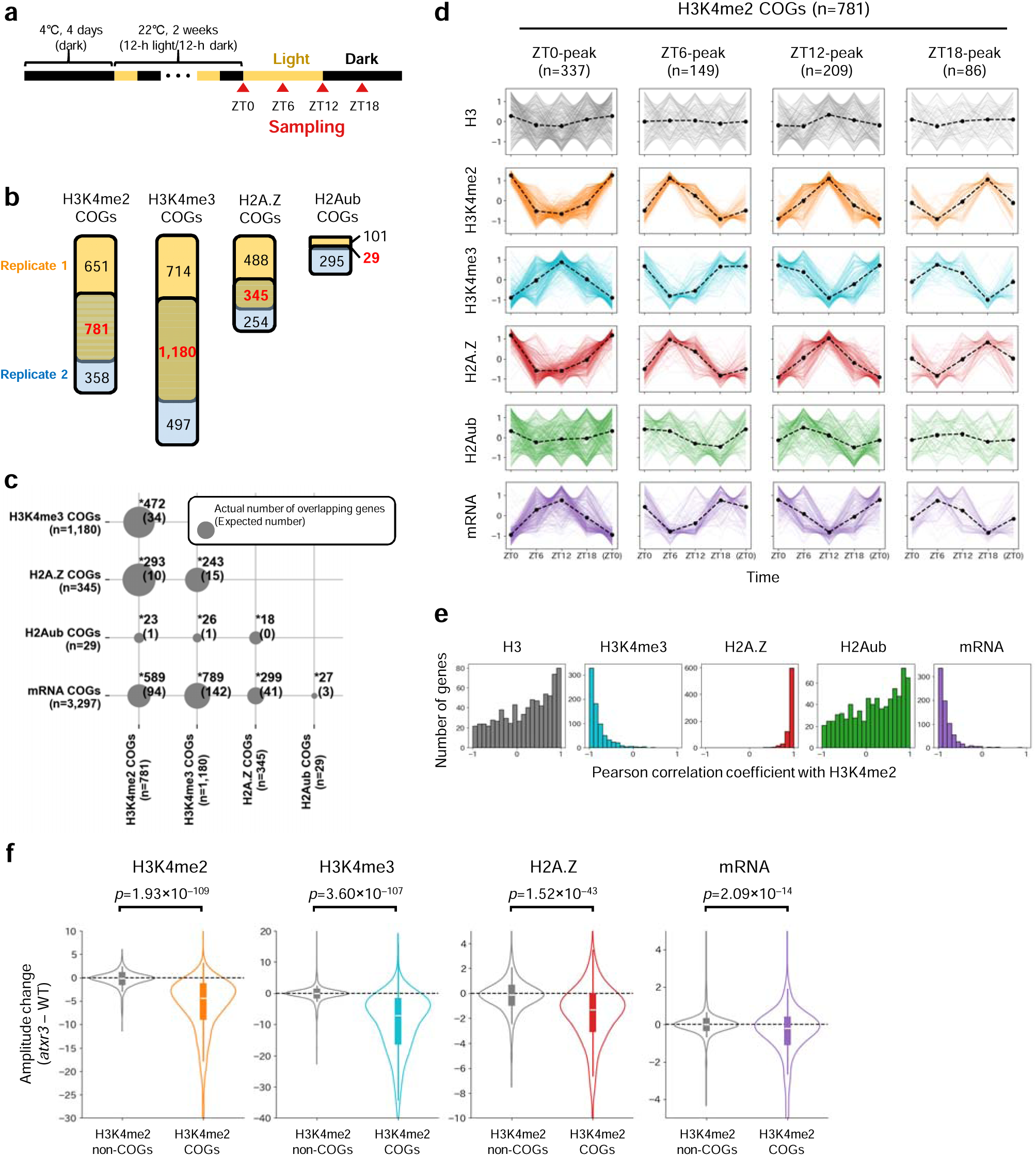
Circadian dynamics of H3K4me2, H3K4me3, H2A.Z, and H2Aub. **a**, Sampling scheme for time-course ChIP-seq and mRNA-seq. **b-e**, Analyses of time-course ChIP-seq and mRNA-seq in WT. **b**, Overlap of genes with H3K4me2, H3K4me3, H2A.Z, and H2Aub oscillation between two biological replicates. These overlapping genes are defined as “H3K4me2 circadian oscillation genes (COGs)” (n=781), “H3K4me3 COGs” (n=1,180), “H2A.Z COGs” (n=345), and “H2Aub COGs” (n=29), respectively. **c**, Overlap between H3K4me2 COGs, H3K4me3 COGs, H2A.Z COGs, H2Aub COGs, and mRNA COGs. Numbers at grid intersections indicate the actual number of overlapping genes; numbers in parentheses indicate the expected number of overlapping genes if the same number of genes is randomly chosen from all protein-coding genes (n=27,443). The size of the circles indicates the similarity (Jaccard index) of the two gene sets. Asterisks indicate significant enrichment (Bonferroni-corrected p-value from one-sided Fisher’s exact test < 0.05). **d**, Circadian changes in H3, H3K4me2, H3K4me3, H2A.Z, H2Aub, and mRNA for H3K4me2 COGs (n=781, as shown in **b**). H3K4me2 COGs are classified into 4 groups based on the time when H3K4me2 reached the maximum (ZT0-peak, ZT6-peak, ZT12-peak, and ZT18-peak). Chromatin mark levels and mRNA levels are standardized to a mean of 0 and a standard deviation of 1 across four time points for each gene. **e**, Histograms showing correlations of temporal changes between H3K4me2 and others (H3, H3K4me3, H2A.Z, H2Aub, and mRNA) for H3K4me2 COGs (n=781). **f**, Violin plots showing the amplitude change (*atxr3* – WT) of H3K4me2, H3K4me3, H2A.Z, and mRNA in H3K4me2 COGs (n=781) and H3K4me2 non-circadian oscillation genes (H3K4me2 non-COGs, 781 genes are randomly selected from genes other than H3K4me2 COGs). Amplitudes are calculated by the difference between the maximum and the minimum values of each chromatin mark level across four time points for each gene. The p-values from two-sided Mann-Whitney U tests are shown.

### The attenuation of H3K4me2 oscillation induces the mirrored attenuation of H2A.Z oscillation

H3K4me2 affects the localization of H2A.Z but not vice versa (Fig. 2, Fig. 3, Fig. 4e, f). Therefore, we speculated that the diurnal oscillation of H3K4me2 also regulates that of H2A.Z. An H3K4 methyltransferase, ARABIDOPSIS TRITHORAX-RELATED3 (ATXR3)/SET DOMAIN GROUP 2 (SDG2), has been reported to regulate the oscillation of H3K4me3 in core clock genes (e.g., *CCA1, LHY* and *TOC1*)^6,31^. We noticed that H3K4me2 COGs were enriched in genes with decreased H3K4me3 in *atxr3* (Supplementary Fig. 8). These findings imply that ATXR3 is involved in the diurnal oscillation of H3K4me2/3. Therefore, we performed time-course ChIP-seq and mRNA-seq using *atxr3* mutants to change the diurnal oscillation pattern of H3K4me2 and investigate the effect on H2A.Z oscillation. Consistent with previous reports, the oscillation of H3K4me3 in most core clock genes was attenuated in *atxr3* (Supplementary Fig. 9). In addition to H3K4me3, H3K4me2 oscillation was weakened (Supplementary Fig. 9), likely reflecting the contribution of ATXR3 on H3K4me2 oscillation^9,32^. Importantly, while H3K4me3 and H3K4me2 oscillate in antiphase in WT, loss of H3K4me3 in *atxr3* did not induce accumulation of H3K4me2 (Supplementary Fig. 9). On the contrary, H3K4me2 oscillation was attenuated in *atxr3* (Fig. 5f; Supplementary Fig. 10a, b). Furthermore, the oscillation of H2A.Z and mRNA was significantly attenuated in *atxr3* (Fig. 5f; Supplementary Fig. 10c, d). The loss of H2A.Z in *atxr3* showed a higher genome-wide correlation with that of H3K4me2 at the same time than that of H3K4me3 in H3K4me2 COGs (Supplementally Fig. 10e), indicating that H2A.Z oscillation is more strongly linked to H3K4me2 oscillation than H3K4me3 oscillation.

In contrast to *atxr3*, approximately 25% of H3K4me2 COGs overlapped with LDL3-target genes, which did not represent significant enrichment (Supplementary Fig. 11a). We also performed time-course ChIP-seq using the *ldl3* mutants. Focusing on H3K4me2 COGs regulated by LDL3, the attenuation of H3K4me2 diurnal oscillation was not observed in *ldl3* partially because both the maximum and the minimum H3K4me2 levels across the four time points significantly increased (Supplementary Fig. 11b). However, the *ldl3* mutation led to changes in the spatiotemporal localization pattern of H3K4me2 in some core clock genes (e.g., *LHY*, *CCA1*, and *TOC1*), which in turn induced similar alterations in H2A.Z (Supplementary Fig. 11c). These results further strengthened the conclusion that H3K4me2 promotes the accumulation of H2A.Z not only in the static state but also in the dynamic state, such as circadian oscillation.

## Discussion

Through genetic analyses, here we demonstrated that H3K4me2 plays a role in regulating the localization of H2A.Z and Polycomb repressive marks H2Aub and H3K27me3 (Fig. 2, Fig. 3). We also showed that H3K4me2 oscillates in a circadian manner even in genes other than core clock genes. Interestingly, H3K4me2 oscillated in phase with H2A.Z but in antiphase with H3K4me3 and mRNA (Fig. 5a-e). This result aligns with our observations in the static state and suggests that H3K4me2, rather than H3K4me3, influences H2A.Z dynamics. Furthermore, the attenuation of H3K4me2 oscillations in the *atxr3* mutant also reduced H2A.Z oscillations (Fig. 5f). Additionally, in the *ldl3* mutant, H2A.Z oscillated while exhibiting an altered intragenic pattern similar to that of H3K4me2 in core clock genes (Supplementary Fig. 11). These results suggest that H3K4me2 has a strong impact on regulating H2A.Z.

Furthermore, despite the modest changes in H2A.Z levels in the mutants with altered H3K4me2 levels, H2A.Z exhibited strong co-oscillation with H3K4me2. This result suggests not only that H3K4me2 regulates H2A.Z but also that shared regulatory factors may coordinate the regulation of both H3K4me2 and H2A.Z. The SWR1-C has been reported to be recruited to core clock genes by the Evening Complex, a key component of the Arabidopsis circadian clock^33^. These components may also recruit factors responsible for H3K4me2 methylation or demethylation.

Although our mutant analysis indicated that H2A.Z, H2Aub, and H3K27me3 act downstream of H3K4me2, only H2A.Z oscillated synchronously with H3K4me2, whereas H2Aub did not (H3K27me3 was reported to lack oscillation^6^). This observation is especially noteworthy considering that overall distribution of H2Aub matches very well to that of H2A.Z (Fig. 1a, Supplementary Fig. 1b). In addition, this observation suggests that the changes in H2A.Z occur independently of Polycomb repressive marks, which is consistent with our finding that H2A.Z levels remained unchanged in the PRC1 and PRC2 mutants (Supplementary Fig. 4, Supplementary Fig. 6). Conversely, alterations in H2A.Z levels affected H3K27me3^18^, suggesting that H2A.Z can regulate Polycomb repressive marks. Additionally, our results suggest a difference in the responsiveness of H2A.Z and H2Aub/H3K27me3. Specifically, the H3K4me2-dependent accumulation of H2A.Z is likely to occur more rapidly than that of H2Aub and H3K27me3. Further studies are necessary to elucidate the molecular mechanisms underlying the circadian oscillations of H3K4me2 and other chromatin marks.

We previously showed that LDL3 demethylates H3K4me2 on actively transcribed genes in a transcription-coupled manner^5^. Based on this and our current findings, we propose a hypothetical model in which H3K4me2 and transcription negatively regulate each other (Fig. 6). In actively transcribed genes, LDL3 cotranscriptionally removes H3K4me2. In contrast, in genes with low transcription activity, H3K4me2 accumulates in the gene body, followed by the relatively rapid accumulation of H2A.Z. Over time, H2Aub and H3K27me3 also accumulate, leading to gene repression. The molecular mechanism by which H3K4me2 recruits downstream chromatin marks remains unknown, but ALFIN1-LIKE (AL) family proteins can be candidates, as they have been shown to bind not only to H3K4me3 but also to H3K4me2 *in vitro*^34,35^. Some AL members bind to ARP6, a component of SWR1-C, which incorporates H2A.Z into nucleosomes^10,11,36,37^. In addition, some AL proteins also bind to PRC1 components^10,38^. AL proteins are conserved in plants but are absent in both animals and fungi, which aligns with the idea that H3K4me2 negatively regulates transcription by promoting the deposition of H2A.Z and H2Aub in a plant-specific manner^34^. Further studies are needed to clarify the molecular mechanism by which H3K4me2 promotes the deposition of H2A.Z, H2Aub, and H3K27me3.

**Fig. 6.**
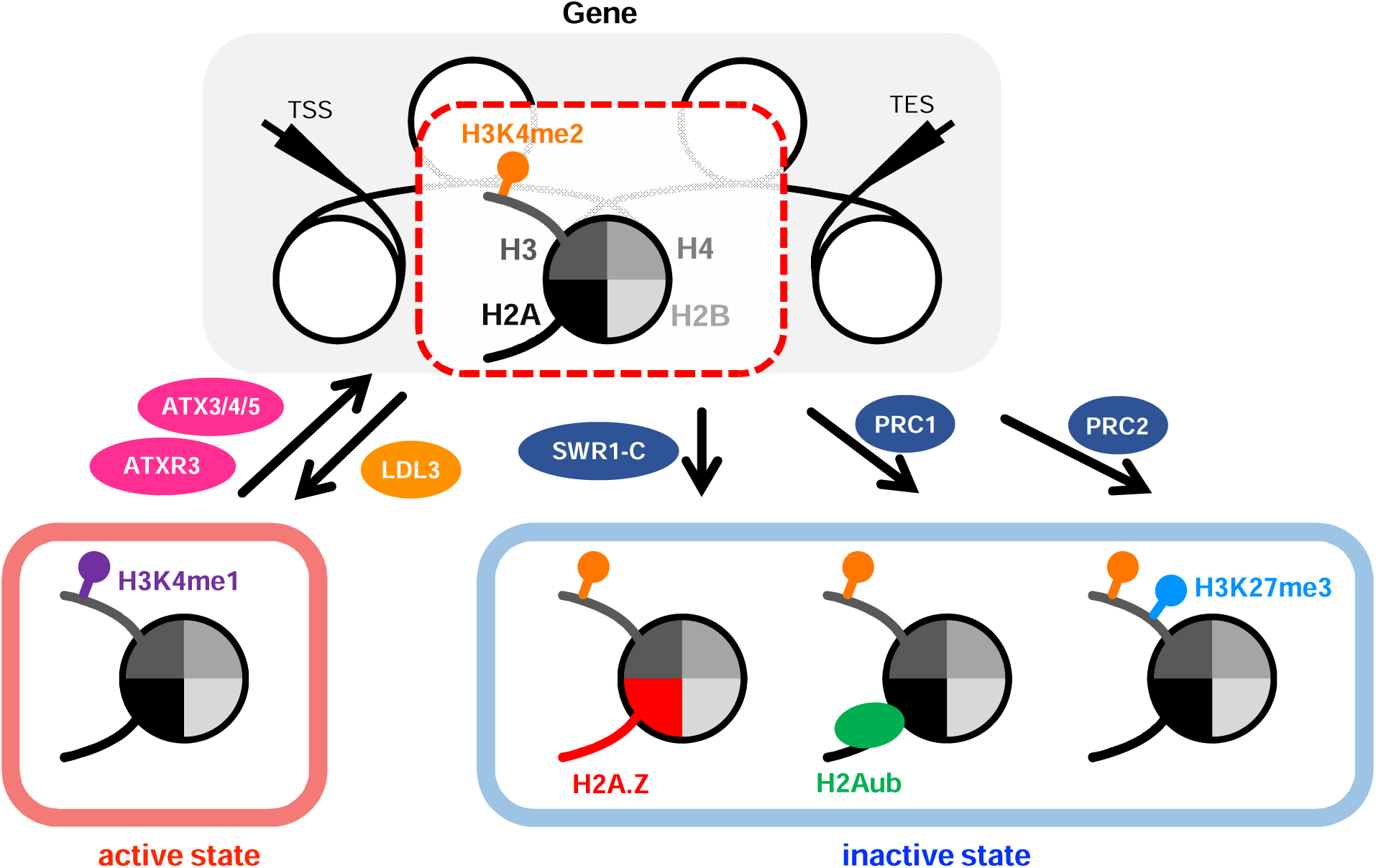
Proposed model in which H3K4me2 regulates H2A.Z, H2Aub and H3K27me3. On highly transcribed genes, LDL3 demethylates H3K4me2 to H3K4me1 in a transcription-coupled manner. On lowly transcribed genes, H3K4me2 recruits SWR1 chromatin remodeling complex (SWR1-C) to deposit H2A.Z, followed by recruiting PRC1 and PRC2 to deposit H2Aub and H3K27me3, respectively.

Although H3K4 methylation is highly conserved across eukaryotes, our findings suggest that its downstream transcriptional regulatory mechanisms have diverged between plants and animals. These results highlight the importance of H3K4me2 in regulating chromatin states and raise intriguing questions about how H3K4me2 influences its downstream long-term chromatin marks, which is conserved from animals but in plants used to memorize environmental stimuli.

## Materials and Methods

### Plant materials and growth conditions

*A. thaliana* strain Columbia-0 (Col-0) was used as wild type (WT). *atx3-1* (GK-128H01), *atx4* (SALK_060156), *atx5* (SAIL_705_H05) and *ldl3-1* (GABI_092C03) were previously described^5,9^. The *jmj32-1*, *elf6-3*, *ref6-1*, *jmj30-2,* and *jmj13G* were previously described^23^. Other mutants used in this paper were described in the following previous reports: *hta9-1* (SALK_054814), *hta11-2* (SALK_031471)^39^, *clf-28* (NASC: N639371)^40^, *bmi1a-1* (SALK_145041), *bmi1b* (WiscDsLox437G06)^27^, *bmi1c-1* (SALK_148143)^41^, *arp6-3* (WiscDS_Los289_29), *pie1-3* (SAIL_78_C11)^42^.

For ChIP-seq and mRNA-seq, seeds were sown on Murashige and Skoog (MS) plates and kept in the dark at 4°C for a few days, then plants were grown for 14 days under long-day conditions (8-h dark/16-h light cycle) at 22°C. For ChIP-seq in *atbmi1a/b/c* or time-course ChIP-seq and mRNA-seq, seeds were sown on MS plates and kept in the dark at 4°C for 4 days, then plants were grown for 14 days under a 12-h light/12-h dark cycle at 22°C.

### Library preparation and sequencing for ChIP-seq

The enhanced chromatin immunoprecipitation (eChIP) approach^43^ was adopted in this study. eChIP-seq was performed with slight modifications to our previously described method^5,44^. ∼0.6g of seedlings were sampled, frozen with liquid nitrogen, ground into a powder using mortar and pestle, and cross-linked with 15 mL of Fixation Buffer (phosphate-buffered saline (PBS) containing 1% formaldehyde, 1mM Pefabloc SC (Roche), cOmplete proteinase inhibitor (Roche) and 0.3% Triton X-100) for 10 min at room temperature (RT). To stop the cross-linking reaction, glycine (final concentration: 200 mM) was added and incubated for 5 min at RT. The cell lysate was centrifuged at 5000 *g* for 5 min at 4°C and the supernatant was discarded. The pellet was washed with 4 mL of ice-cold PBS, resuspended with 0.8 mL of Buffer S (50 mM HEPES-KOH (pH7.5), 150 mM NaCl, 1 mM Ethylene Diamine Tetraacetic Acid (EDTA), 1% Triton X-100, 0.1% sodium deoxycholate and 1% Sodium Dodecyl Sulfate (SDS)) containing cOmplete proteinase inhibitor, and incubated for 10 min at 4°C. The chromatin in the cell lysate was fragmented by sonication using S220 Focused-ultrasonicator (Covaris) and milliTUBE 1 ml AFA Fiber (Covaris) with following settings (time: 20-25 min, peak power: 140, duty factor: 5.0, cycles/burst: 200 and avg. power: 7.0). After centrifuging at 20,000 *g* for 3 min at 4°C, the supernatant was discarded and the pellet was resuspended with Buffer F (50 mM HEPES-KOH (pH7.5), 150 mM NaCl, 1 mM EDTA, 1% Triton X-100 and 0.1% sodium deoxycholate) containing cOmplete proteinase inhibitor. For antibody reaction, 1-2 µL of antibodies were added to the chromatin solution, and this mixture was incubated overnight at 4°C. The following antibodies were used; anti-H3 (ab1791, Abcam), anti-H3K4me2 (ab32356, Abcam), anti-H3K4me3 (ab8580, Abcam), anti-H3K27me3 (MABI0323, Cosmo Bio), anti-H2A.Z (ref. ^45^) and anti-H2AK119ub (#8240, Cell Signaling) antibodies. The anti-H2AK119ub antibody was reported to recognize Arabidopsis H2AK121 (K120 excluding the N-terminal methionine)^27^. The chromatin-antibody mixture was incubated with Dynabeads M-280 Sheep anti-Mouse IgG (Thermo Fisher Scientific) for anti-H3K27me3 or Dynabeads Protein G (Thermo Fisher Scientific) for other antibodies for 2 hours at 4°C with rotation. The beads were washed once with low-salt RIPA buffer (50 mM HEPES-KOH (pH7.5), 150 mM NaCl, 1 mM EDTA, 1% Triton X-100, 0.1% sodium deoxycholate and 0.1% SDS) containing cOmplete proteinase inhibitor, twice with medium-salt RIPA buffer (low-salt RIPA buffer with NaCl concentration changed from 150 mM to 300 mM) or high-salt RIPA buffer (from 150 mM to 500 mM) containing cOmplete proteinase inhibitor, once with ChIP wash buffer (10 mM Tris-HCl (pH7.8), 250 mM LiCl, 1% Igepal CA-630, 1 mM EDTA, 1% sodium deoxycholate) and once with TE buffer. The chromatin was detached from the beads by adding ChIP elution buffer (10 mM Tris-HCl (pH 7.8), 300 mM NaCl, 5 mM EDTA, and 0.5% SDS) with 2 µL of RNase (10 mg/ml; NIPPON GENE) and incubating for 30 min at 37°C. Then, 2 µL of Proteinase K (20 mg/ml; Thermo Fisher Scientific) was added and incubated for over 3 hours at 65°C. The immunoprecipitated DNA was purified using the Monarch PCR & DNA Cleanup Kit (New England Biolabs). The libraries for ChIP-seq were prepared using ThruPLEX DNA-Seq Kit (Takara Bio) and DNA Unique Dual Index Kit (Takara Bio) following the manufacturer’s protocol and purified using SPRIselect beads (Beckman Coulter). The libraries were sequenced by the HiSeq X or NovaSeq X Plus sequencer (Illumina). Two biological replicates were analyzed using independently grown plants.

For time-course ChIP-seq, 0.6 g of seedlings were sampled at ZT0, ZT6, ZT12, and ZT18 (Fig. 5a). The procedure of ChIP-seq was the same as above.

### Library preparation and sequencing for mRNA-seq

Three seedlings were sampled at ZT0, ZT6, ZT12, and ZT18 (Fig. 5a) for three biological replicates. A seedling was frozen with liquid nitrogen and ground into a powder using Automill (Token) with two stainless steel beads at 1,600 r/min for 60 sec. Total RNA was purified using RNeasy Plant Mini Kit (Qiagen). The libraries for mRNA-seq were prepared using KAPA RNA HyperPrep Kit (Roche) following the manufacturer’s protocol and sequenced by NovaSeq X Plus sequencer (Illumina).

### Sequence data processing

The sequenced data of ChIP-seq was trimmed of adaptors using Trimmomatic (v.0.39)^46^, then mapped to the TAIR10 reference genome using Bowtie2 (v.2.4.5)^47^. Multi-mapped reads were removed from mapped reads, and the number of reads on each gene was counted using the “coverage” command of BEDtools (v.2.30.0)^48^.

The sequenced data of mRNA-seq was trimmed of adaptors using Trimmomatic (v.0.39)^46^, then mapped to the TAIR10 reference genome using STAR (v.2.7.9a)^49^. The number of reads on each gene was counted using the “coverage” command of BEDtools (v.2.30.0)^48^.

Heatmaps and metaplots were generated using deepTools (v.3.5.1)^50^. Genome browser views of ChIP-seq signals were displayed with Integrative Genomics Viewer (v.2.14.1)^51^. Other figures were generated using Python libraries: pandas (v.2.2.3), matplotlib (v.3.3.4), and seaborn (v.0.13.0). For the detection of H2A.Z peaks against H3, MACS2 (v2.2.7.1) with default parameters was used (Supplementary Fig. 5a)^52^.

### Statistical analysis

The changes in gene expression levels between WT and *ldl3* (Supplementary Fig. 2d) and WT and *atx3/4/5* (Supplementary Fig. 3d) were tested with Wald test using an R package DESeq2 (v.1.34.0)^53^. The enrichment between H3K4me2 COGs, H3K4me3 COGs, H2A.Z COGs, H2Aub COGs, and mRNA COGs was evaluated with one-sided Fisher’s exact test with Bonferroni-correction using a Python library SciPy (v.1.12.0) (Fig. 5c). The enrichment between H3K4me2 COGs and LDL3-target genes was evaluated with one-sided Fisher’s exact test using SciPy (v.1.12.0) (Supplementary Fig. 11a). Two-sided Mann-Whitney U tests were performed using SciPy (v.1.12.0) for evaluating the differences between two gene groups (Fig. 2c, Fig. 3c, Fig. 5f; Supplementary Fig. 8b, Supplementary Fig. 11b).

### Definition of circadian oscillating genes (COGs) of chromatin marks and mRNA

H3K4me2 COGs were selected with the following 2 criteria; When comparing any two time points (e.g., ZT0 and ZT6), (1) There was a genomic region within that gene where H3K4me2 was significantly enriched at one time point compared to another time point (likelihood ratio ≥ 1000), and (2) the H3K4me2 level (RPKM) in that gene at one time was 1.2 times higher than that at another time point. As for criterion 1, the “bdgdiff” command of MACS3 (v.3.0.1) was used to detect H3K4me2-enriched regions^52^. H3K4me3 COGs, H2A.Z COGs, and H2Aub COGs were also selected similarly.

mRNA COGs were selected with the following three criteria: (1) the expression level significantly changed across four time points (ZT0, ZT6, ZT12, and ZT18), with the adjusted p-value of the likelihood ratio test across four time points being < 10^−5^, (2) the mean expression (RPKM) across four time points was > 2, and (3) the fold change (max/min) was > 1.5. The likelihood ratio test was performed using DESeq2 (v.1.34.0)^53^. Two independent biological replicates were analyzed in parallel and the selected genes in both replicates were defined as mRNA COGs.

### Sources of reanalyzed data

The following data was reanalyzed in this study: mRNA-seq in WT and *ldl3* (NCBI: PRJNA934737)^5^ and ChIP-seq for H3K4me1, H3K4me2, and H3K4me3 in WT and *atxr3* (NCBI: PRJNA732996)^9^.

## Data availability

The data have been deposited with links to BioProject accession number PRJNAXXXXXXX in the NIH BioProject database.

## Acknowledgements

The computations were partially performed on the NIG supercomputer at NIG, Japan. We thank Taiko To for kindly providing H2A.Z antibodies. We thank Nobutoshi Yamaguchi, Taku Sasaki, Taiko To, and Akihisa Osakabe for kindly providing mutant seeds. This work was supported by grants from Human Frontier Science Program (HFSP) (RGP0025/2021) to T.K., Japan Science and Technology Agency (JST) CREST (no. JPMJCR15O1) to H.K. and T.K., Japan Society for the Promotion of Science (JSPS) (nos. 21H04977 to H.K. and T.K., 23H00365 and 24K21268 to T.K., and 20H05913, 23K23565, and 25K02252 to S.I.).

## Author Contributions

T.N., S.M., S.O., H.N., H.K., S.I., and T.K. conceived the study. T.N., S.M., and S.O. performed the experiments. T.N. conducted the analysis and made figures. T.N. drafted and T.N., S.M., H.N., H.K., S.I., and T.K. edited the manuscript.

---

**Supplementary Fig. 1 | H3K4me2 is more positively correlated with H2A.Z, H2Aub, and H3K27me3 than H3K4me1 and H3K4me3.**

**a**, The comparison between the level of H3K4 methylation (H3K4me1/2/3) and H3K27me3, H2A.Z, and H2Aub. **b**, The comparison between the level of H2A.Z, H2Aub, and H3K27me3. Each dot represents the square root of RPKM within protein-coding genes (n=27,443). ρ: Spearman’s rank correlation coefficient.

**Supplementary Fig. 2 | ChIP-seq results that H2A.Z and H2Aub increase in *ldl3* are reproducible.**

**a**, Scatter plots showing the levels of H3K4me2, H2A.Z, H2Aub, and H3K27me3 comparing WT and *ldl3*. Each dot represents a protein-coding gene. Red dots indicate LDL3-target genes (n=7,115, as shown in Fig. 2a). **b**, Metaplots showing the averaged distribution of H3K4me2, H2A.Z, H2Aub, and H3K27me3 in WT (black) and *ldl3* (red) in LDL3-target genes (n=7,115) and all protein-coding genes (n=27,443). **c,** Box plots showing the relationship between the change in H3K4me2 and the change in H2A.Z, H2Aub, and H3K27me3 in *ldl3* compared to WT. Histograms above the boxplots indicate the number of genes in each category of H3K4me2 change. The values of scatter plots, metaplots, and box plots represent RPKM. **d**, The expression levels of H3K27me3-related, H2A.Z-related, and H2Aub-related genes in WT (black, n=3) and *ldl3* (red, n=2). The previous RNA-seq data was used^5^. The values represent the square root of RPKM. The differences in gene expression levels between WT and *ldl3* were tested by Wald test with DESeq2. n.s.: not significant (The p-value from Wald test adjusted by Benjamini-Hochberg correction > 0.05). N/A: The gene was filtered out due to the low expression level when calculating the adjusted p-value.

**Supplementary Fig. 3 | ChIP-seq results that H2A.Z, H2Aub, and H3K27me3 decrease in *atx3/4/5* are reproducible.**

**a**, Scatter plots showing the levels of H3K4me2, H2A.Z, H2Aub and H3K27me3 comparing WT and *atx3/4/5*. Each dot represents a protein-coding gene. Blue dots indicate ATX3/4/5-target genes (n=5,842, as shown in Fig. 3a). **b,** Metaplots showing the averaged distribution of H3K4me2, H2A.Z, H2Aub, and H3K27me3 in WT (black) and *atx3/4/5* (blue) in ATX3/4/5-target genes (n=5,842) and all protein-coding genes (n=27,743). **c,** Box plots showing the relationship between the change in H3K4me2 and the change in H2A.Z, H2Aub, and H3K27me3 in *atx3/4/5* compared to WT. Histograms above the boxplots indicate the number of genes in each category of H3K4me2 change. The values of scatter plots, metaplots, and box plots represent RPKM. **d**, The expression levels of H3K27me3-related, H2A.Z-related and H2Aub-related genes in WT (black, n=3) and *atx3/4/5* (blue, n=3). The values represent the square root of RPKM. The differences in gene expression levels between WT and *atx3/4/5* were tested by Wald test with DESeq2. n.s.: not significant (The p-value from Wald test adjusted by Benjamini-Hochberg correction > 0.05). N/A: The gene was filtered out due to the low expression level when calculating the adjusted p-value.

**Supplementary Fig. 4 | ChIP-seq for H3K27me3 and H2A.Z in *jmj30 jmj32 elf6 ref6 jmj13* quintuple mutants (*jmj* quintuple) and *clf*.**

**a**, The overlap of genes with increased H3K27me3 in *jmj* quintuple compared to WT (H3K27me3 change (*jmj* quintuple–WT) > 5) between two biological replicates. JMJ-target genes are defined as the overlapping genes (n=3,048). **b**, The overlap of genes with decreased H3K27me3 in *clf* compared to WT (H3K27me3 change (*clf*–WT) < –5) between two biological replicates. CLF-target genes are defined as the overlapping genes (n=2,160). **c**, Scatter plots showing the levels of H2A.Z comparing WT and *jmj* quintuple. Each dot represents a protein-coding gene. Red dots indicate JMJ-target genes (n=3,048). **d**, Scatter plots showing the levels of H2A.Z comparing WT and *clf*. Each dot represents a protein-coding gene. Blue dots indicate CLF-target genes (n=2,160). **e**, Metaplots showing the distribution of H2A.Z in JMJ-target genes (n=3,048) in WT and *jmj* quintuple. **f**, Metaplots showing the distribution of H2A.Z in CLF-target genes (n=2,160) in WT and *clf*. The values of scatter plots and metaplots represent RPKM.

**Supplementary Fig. 5 | ChIP-seq for H3K4me2 and H2A.Z in *arp6* and *pie1*. a**, Positions of H2A.Z peak summits relative to TSS (x-axis) and frequency of H2A.Z peak (y-axis) in WT. H2A.Z peak summits were detected against H3 using MACS2 peak caller. The genic regions from TSS to TSS+500bp (TSS+500 region) were analyzed. **b**, Scatter plots comparing H2A.Z changes in *arp6* and *pie1*. Each dot represents a protein-coding gene. **c**, Venn diagram showing the overlap between genes with decreased H2A.Z in *arp6* (H2A.Z change (*arp6*–WT) < –15) and genes with decreased H2A.Z in *pie1* (H2A.Z change (*pie1*–WT) < –15). SWR1-C-target genes are defined as the overlapping genes (n=6,096). **d**, Scatter plots showing the levels of H2A.Z (left) and H3K4me2 (right) comparing WT and *arp6*. Each dot represents a protein-coding gene. Blue dots indicate SWR1-C-target genes (n=6,096). **e**, Scatter plots showing the relationship between H2A.Z changes and H3K4me2 changes in *arp6* (left) and *pie1* (right) compared to WT. The values of scatter plots represent the RPKM of TSS+500 regions. ρ: Spearman’s rank correlation coefficient.

**Supplementary Fig. 6 | ChIP-seq for H2Aub and H2A.Z in *bmi1a/b/c*.**

**a**, The overlap of genes with decreased H2Aub in *atbmi1a/b/c* (H2Aub change (*atbmi1a/b/c*–WT) < –2) between two biological replicates. AtBMI1-target genes are defined as the overlapping genes (n=4,982). **b,** Scatter plots showing H2A.Z levels in *atbmi1a/b/c* compared to WT. Each dot represents a protein-coding gene. Blue dots indicate AtBMI1-target genes (n=4,982). **c,** Metaplots show the averaged distribution of H2A.Z in AtBMI1-target genes (n=4,982) in WT and *atbmi1a/b/c*. The values of scatter plots and metaplots represent RPKM.

**Supplementary Fig. 7 | Chromatin mark and mRNA dynamics at core clock genes in WT.**

**a**, Temporal changes in H3K4me2 (orange), H3K4me3 (cyan), H2A.Z (red), H2Aub (green), and mRNA (purple) at core clock genes. The first y-axis shows the levels of chromatin marks normalized by H3. The second y-axis shows the levels of gene expression (RPKM). For ChIP-seq data, dots and triangles indicate replicate 1 and 2, respectively, and lines indicate their mean. For mRNA-seq data, three biological replicates were used. Dots indicate the mean of them, and error bars indicate standard error. **b**, Genome browser views showing the temporal changes of H3K4me2 (orange), H3K4me3 (cyan), H2A.Z (red) and H2Aub (green) at *LHY*, *CCA1*, and *TOC1*.

**Supplementally Fig. 8 | H3K4me2 circadian oscillation genes (COGs) are enriched in ATXR3-target genes.**

**a-b**, Reanalysis of the previous ChIP-seq data for H3K4 methylation in WT and *atxr3*^9^. **a**, Scatter plots showing the levels of H3K4me1, H3K4me2, and H3K4me3 in *atxr3* compared to WT. Each dot represents a protein-coding gene. Red dots represent H3K4me2 COGs (n=781, as shown in Fig. 5b). **b**, Box plots showing the changes of H3K4me1, H3K4me2, and H3K4me3 in *atxr3* compared to WT. Red boxes indicate H3K4me2 COGs (n=781) and grey boxes indicate H3K4me2 non-circadian oscillation genes (H3K4me2 non-COGs, 781 genes are randomly selected from genes other than H3K4me2 COGs). The p-values from two-sided Mann-Whitney U tests are shown. n.s.: not significant (*p* > 0.05).

**Supplementary Fig. 9 | Chromatin marks and mRNA dynamics at core clock genes in WT and *atxr3*.**

Genome browser views showing the temporal changes of H3K4me2 (orange), H3K4me3 (cyan), and H2A.Z (red) at *LHY*, *CCA1*, *TOC1*, *PRR9*, *PRR7,* and *LUX* in WT and *atxr3*. Line plots next to the genome browser views show the level of H3K4me2, H3K4me3, and H2A.Z normalized by H3 in WT (black) and *atxr3* (red).

**Supplementary Fig. 10 | The loss-of-function of ATXR3 extinguishes H3K4me3 oscillation and attenuates H3K4me2, H2A.Z, and mRNA oscillation.**

**a-d**, Scatter plots showing the relationship of H3K4me2 change (**a**), H3K4me3 change (**b**), H2A.Z change (**c**), and mRNA change (**d**) over 6 hours between in WT and that in *atxr3* in H3K4me2 circadian oscillation genes (n=781, as shown in Fig. 5b). Line plots showing linear regression. **e**, Heatmaps showing the Spearman correlation coefficient between the fold change (*atxr3*/WT) of H3K4me2 and H2A.Z (left), and H3K4me3 and H2A.Z (right) in H3K4me2 circadian oscillation genes (n=781) at each time point.

**Supplementary Fig. 11 | Chromatin marks and mRNA dynamics in WT and *ldl3*.**

**a**, The overlap between H3K4me2 circadian oscillation genes (H3K4me2 COGs, as shown in Fig, 5b) and LDL3-target genes (as shown in Fig. 2a). The enrichment of them is not significant (n.s., the p-value from Fisher’s exact test > 0.05). **b**, Violin plots showing the changes in the amplitude, the maximum value, and the minimum value of H3K4me2 diurnal oscillation in *ldl3* compared to WT. Genes that belong to H3K4me2 COGs but not LDL3-target genes (grey, n=582) and genes that belong to both H3K4me2 COGs and LDL3-target genes (red, n=199) are shown. The p-values from two-sided Mann-Whitney U tests are shown. **c**, Genome browser views showing the temporal changes of H3K4me2 (orange), H3K4me3 (cyan), and H2A.Z (red) at *LHY*, *CCA1*, *TOC1*, *PRR5*, *PRR3*, and *GI*. Line plots next to the genome browser views show the level of H3K4me2, H3K4me3, and H2A.Z normalized by H3 in WT (black) and *ldl3* (red).

