## Supplemental Figure for "H3K4me2 orchestrates H2A.Z and Polycomb repressive marks in *Arabidopsis*"

### Slide 1
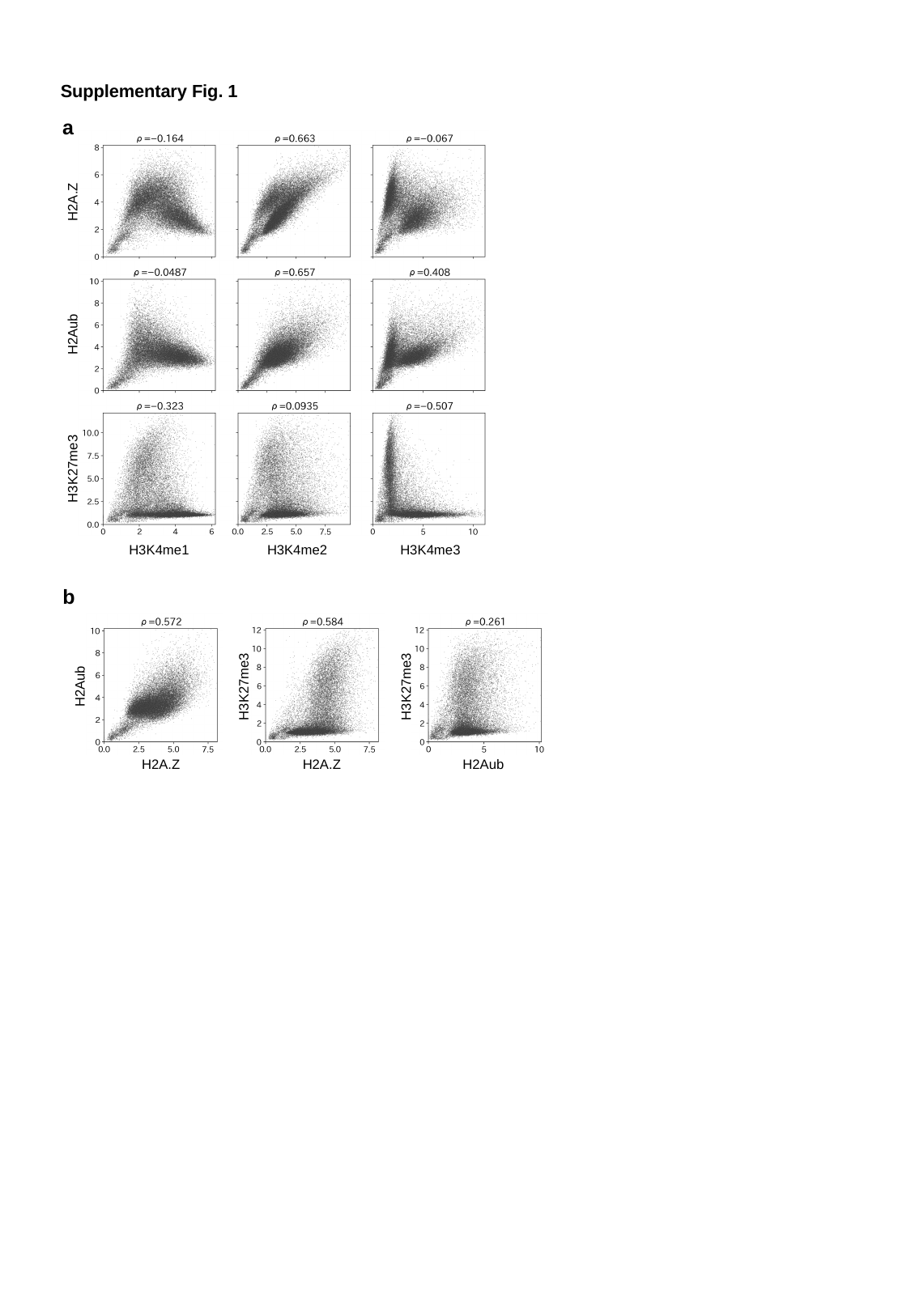

Supplementary Fig. 1
a
H2A.Z
H2Aub
H3K27me3
H3K4me1
H3K4me2
H3K4me3
b
H2Aub
H3K27me3
H3K27me3
H2A.Z
H2A.Z
H2Aub

### Slide 2
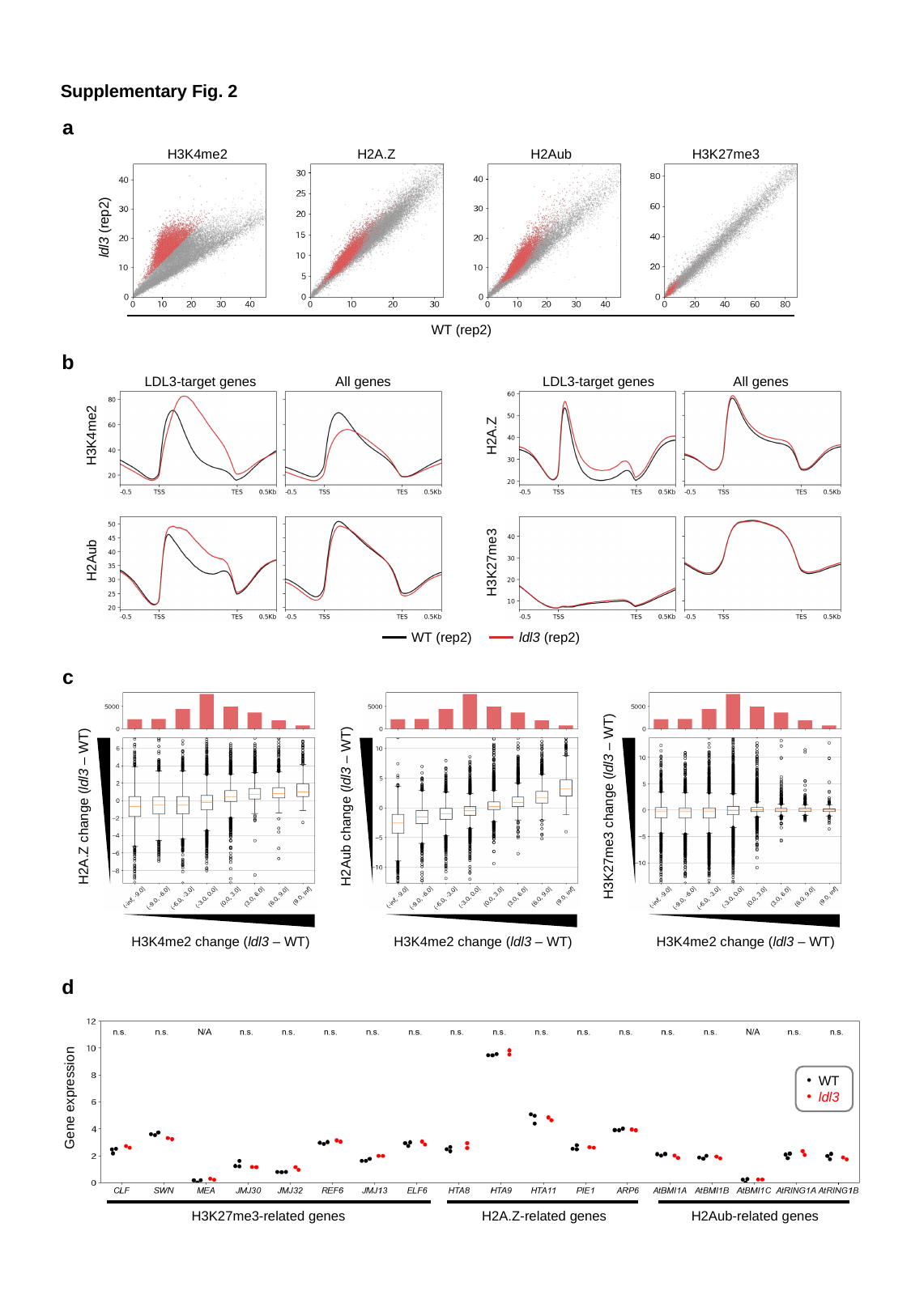

Supplementary Fig. 2
a
H3K4me2
H2A.Z
H2Aub
H3K27me3
ldl3 (rep2)
WT (rep2)
b
LDL3-target genes
All genes
LDL3-target genes
All genes
H3K4me2
H2A.Z
H2Aub
H3K27me3
WT (rep2)
ldl3 (rep2)
c
H3K27me3 change (ldl3 – WT)
H3K4me2 change (ldl3 – WT)
H2Aub change (ldl3 – WT)
H3K4me2 change (ldl3 – WT)
H2A.Z change (ldl3 – WT)
H3K4me2 change (ldl3 – WT)
d
WT
ldl3
Gene expression
H2Aub-related genes
H3K27me3-related genes
H2A.Z-related genes

### Slide 3
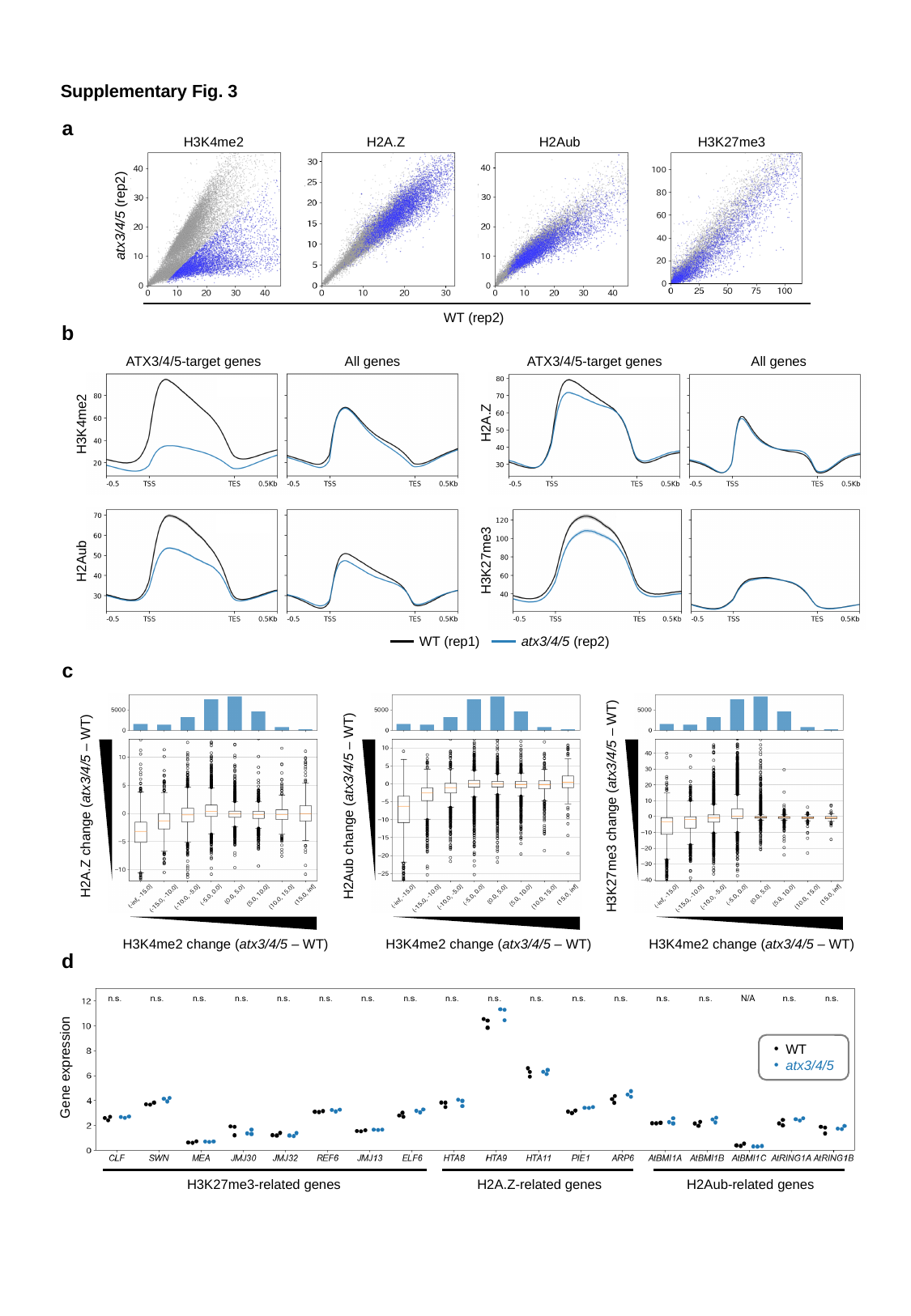

Supplementary Fig. 3
a
H3K4me2
H2A.Z
H2Aub
H3K27me3
atx3/4/5 (rep2)
WT (rep2)
b
ATX3/4/5-target genes
All genes
ATX3/4/5-target genes
All genes
H2A.Z
H3K4me2
H3K27me3
H2Aub
WT (rep1)
atx3/4/5 (rep2)
c
H3K27me3 change (atx3/4/5 – WT)
H3K4me2 change (atx3/4/5 – WT)
H2Aub change (atx3/4/5 – WT)
H3K4me2 change (atx3/4/5 – WT)
H2A.Z change (atx3/4/5 – WT)
H3K4me2 change (atx3/4/5 – WT)
d
WT
atx3/4/5
Gene expression
H2Aub-related genes
H3K27me3-related genes
H2A.Z-related genes

### Slide 4
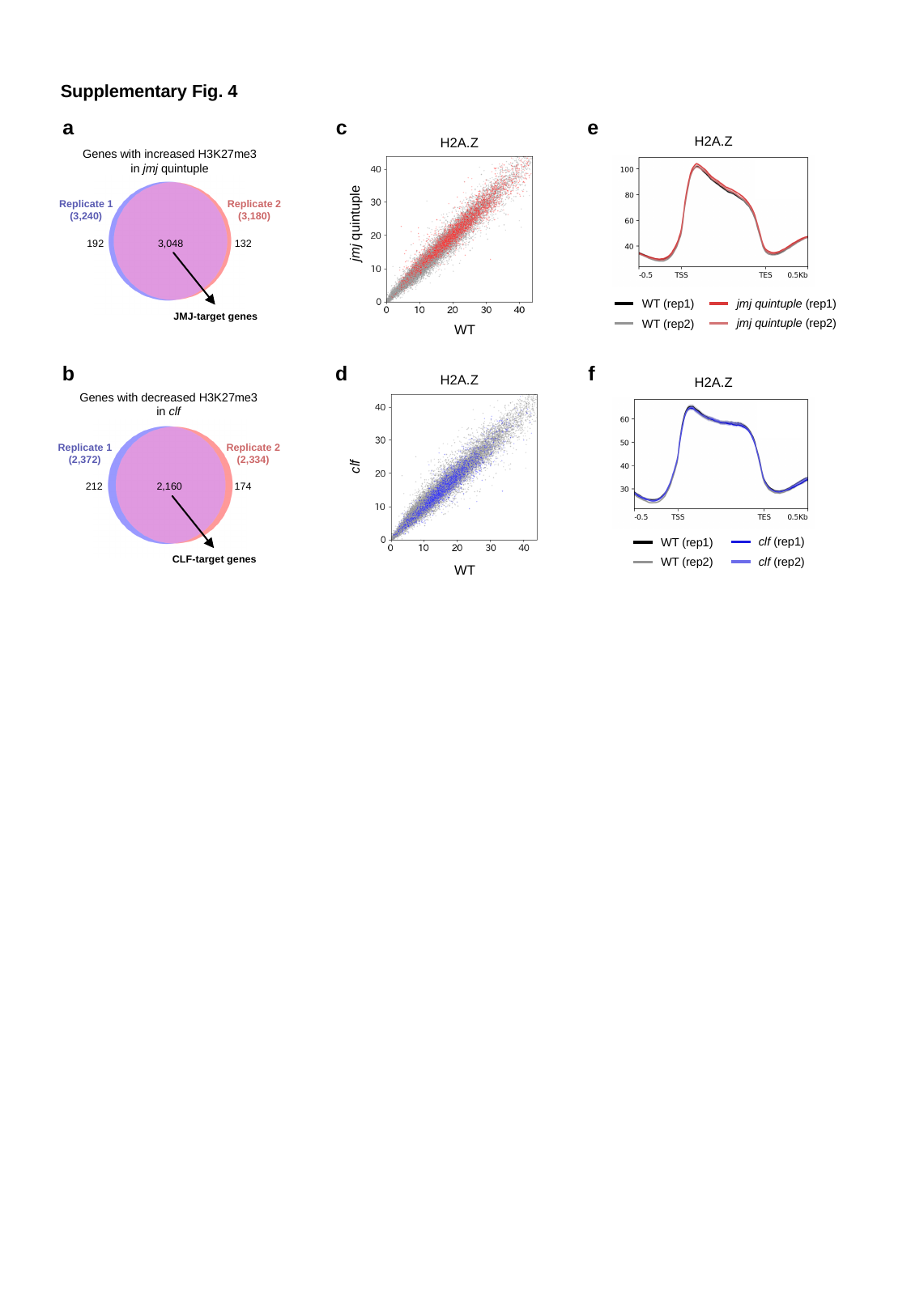

Supplementary Fig. 4
a
c
e
H2A.Z
H2A.Z
Genes with increased H3K27me3
in jmj quintuple
Replicate 1
(3,240)
Replicate 2
(3,180)
192
3,048
132
JMJ-target genes
jmj quintuple
jmj quintuple (rep1)
WT (rep1)
jmj quintuple (rep2)
WT (rep2)
WT
b
d
f
H2A.Z
H2A.Z
Genes with decreased H3K27me3
in clf
Replicate 1
(2,372)
Replicate 2
(2,334)
212
2,160
174
CLF-target genes
clf
clf (rep1)
WT (rep1)
clf (rep2)
WT (rep2)
WT

### Slide 5
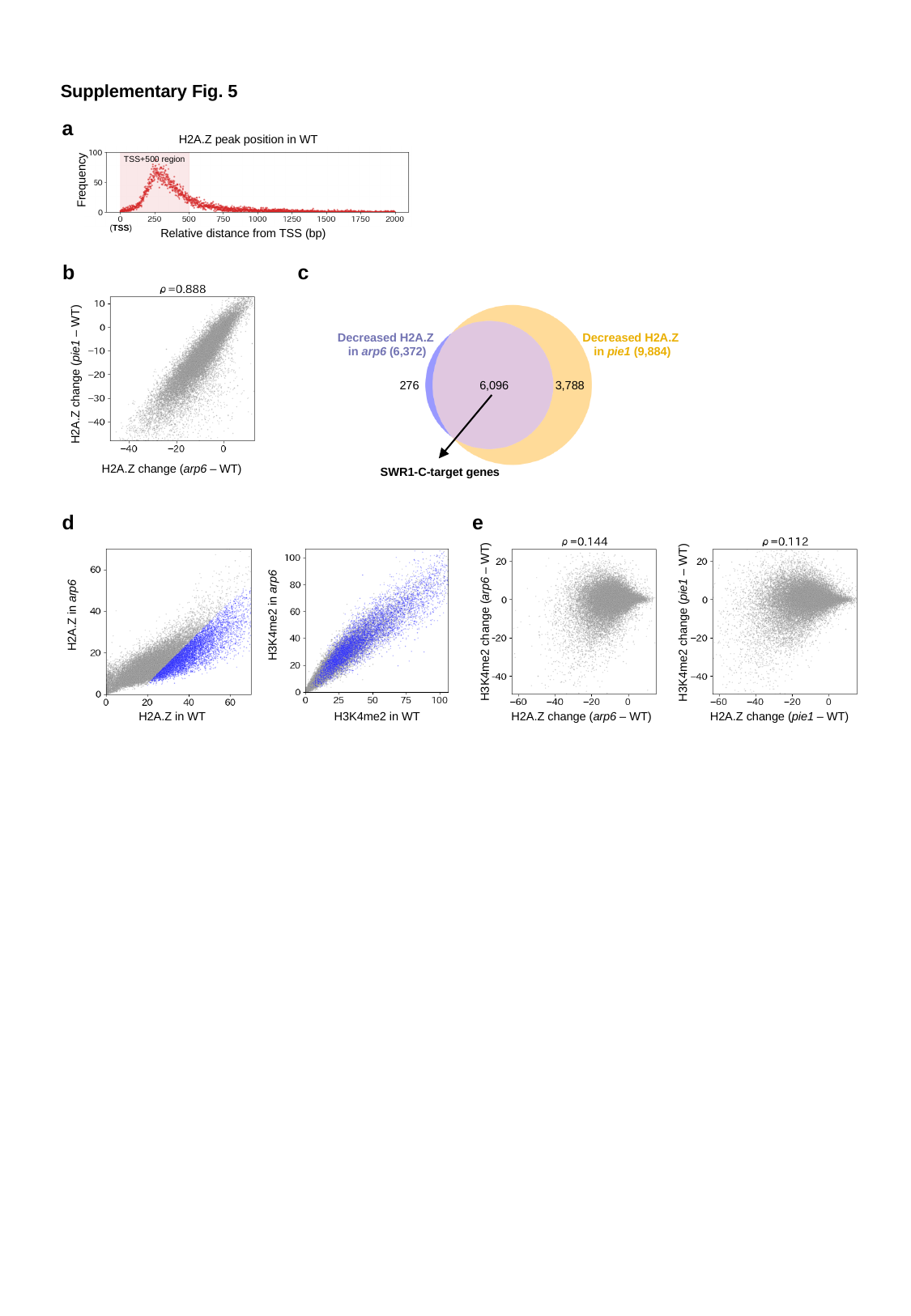

Supplementary Fig. 5
a
H2A.Z peak position in WT
TSS+500 region
Frequency
(TSS)
Relative distance from TSS (bp)
b
c
H2A.Z change (pie1 – WT)
H2A.Z change (arp6 – WT)
Decreased H2A.Z
in arp6 (6,372)
Decreased H2A.Z
 in pie1 (9,884)
276
6,096
3,788
SWR1-C-target genes
d
e
H3K4me2 in arp6
H2A.Z in arp6
H3K4me2 change (arp6 – WT)
H3K4me2 change (pie1 – WT)
H2A.Z in WT
H3K4me2 in WT
H2A.Z change (arp6 – WT)
H2A.Z change (pie1 – WT)

### Slide 6
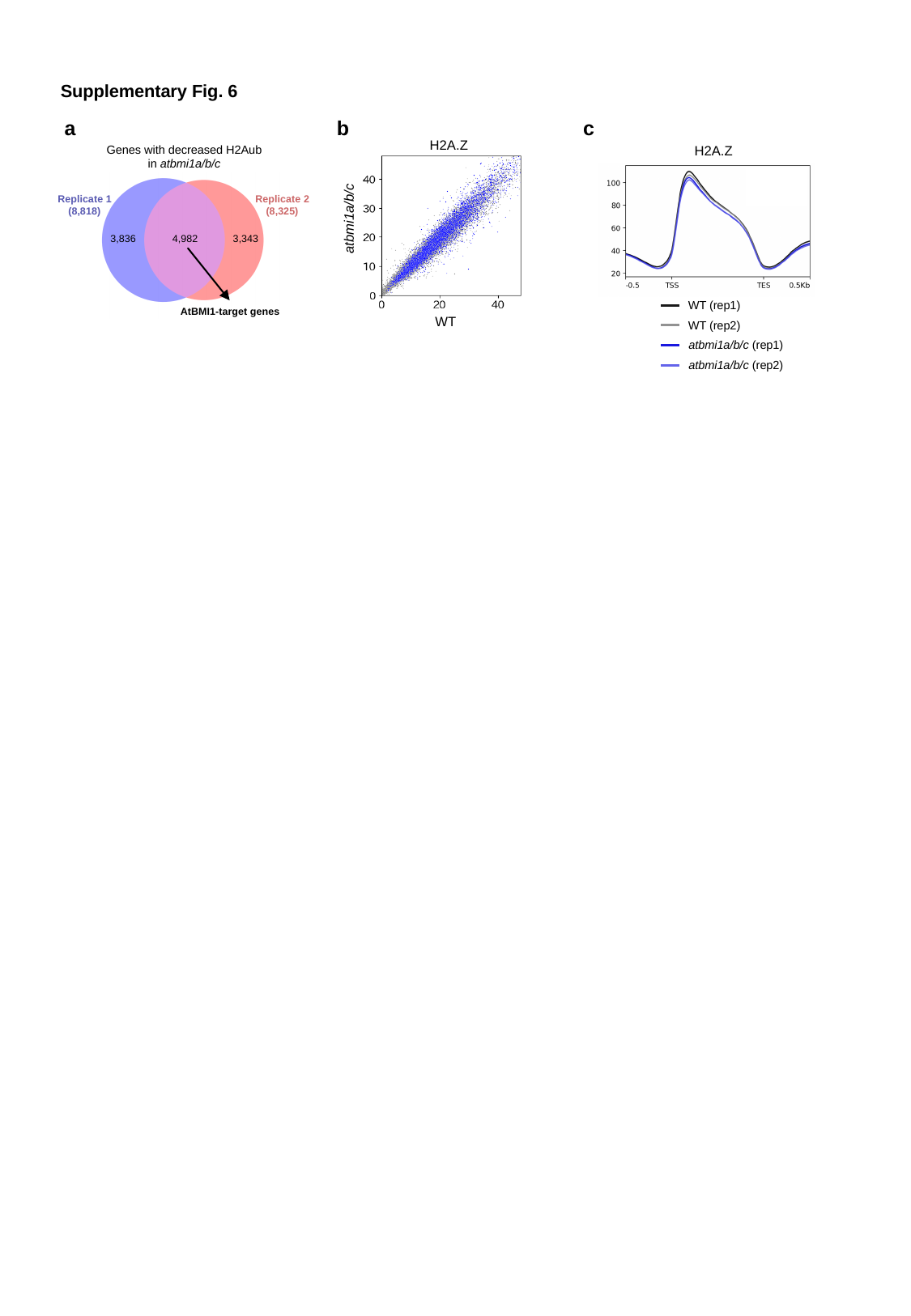

Supplementary Fig. 6
a
b
c
H2A.Z
Genes with decreased H2Aub
in atbmi1a/b/c
Replicate 1
(8,818)
Replicate 2
(8,325)
3,836
4,982
3,343
AtBMI1-target genes
H2A.Z
atbmi1a/b/c
WT (rep1)
WT (rep2)
atbmi1a/b/c (rep1)
atbmi1a/b/c (rep2)
WT

### Slide 7
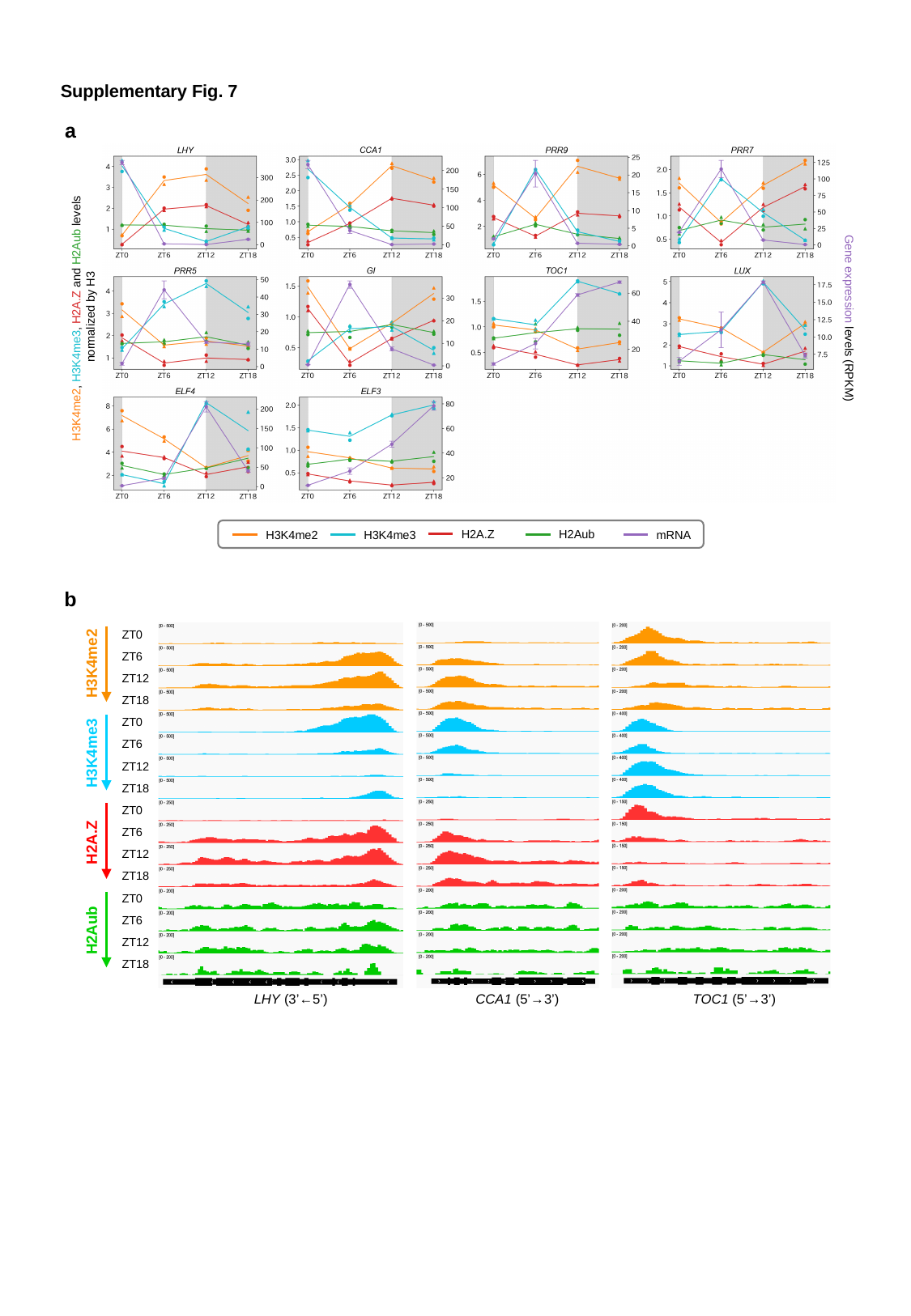

Supplementary Fig. 7
a
H3K4me2, H3K4me3, H2A.Z and H2Aub levels
 normalized by H3
Gene expression levels (RPKM)
H2A.Z
H2Aub
H3K4me3
mRNA
H3K4me2
b
ZT0
ZT6
H3K4me2
ZT12
ZT18
ZT0
ZT6
H3K4me3
ZT12
ZT18
ZT0
ZT6
H2A.Z
ZT12
ZT18
ZT0
ZT6
H2Aub
ZT12
ZT18
LHY (3’←5’)
CCA1 (5’→3’)
TOC1 (5’→3’)

### Slide 8
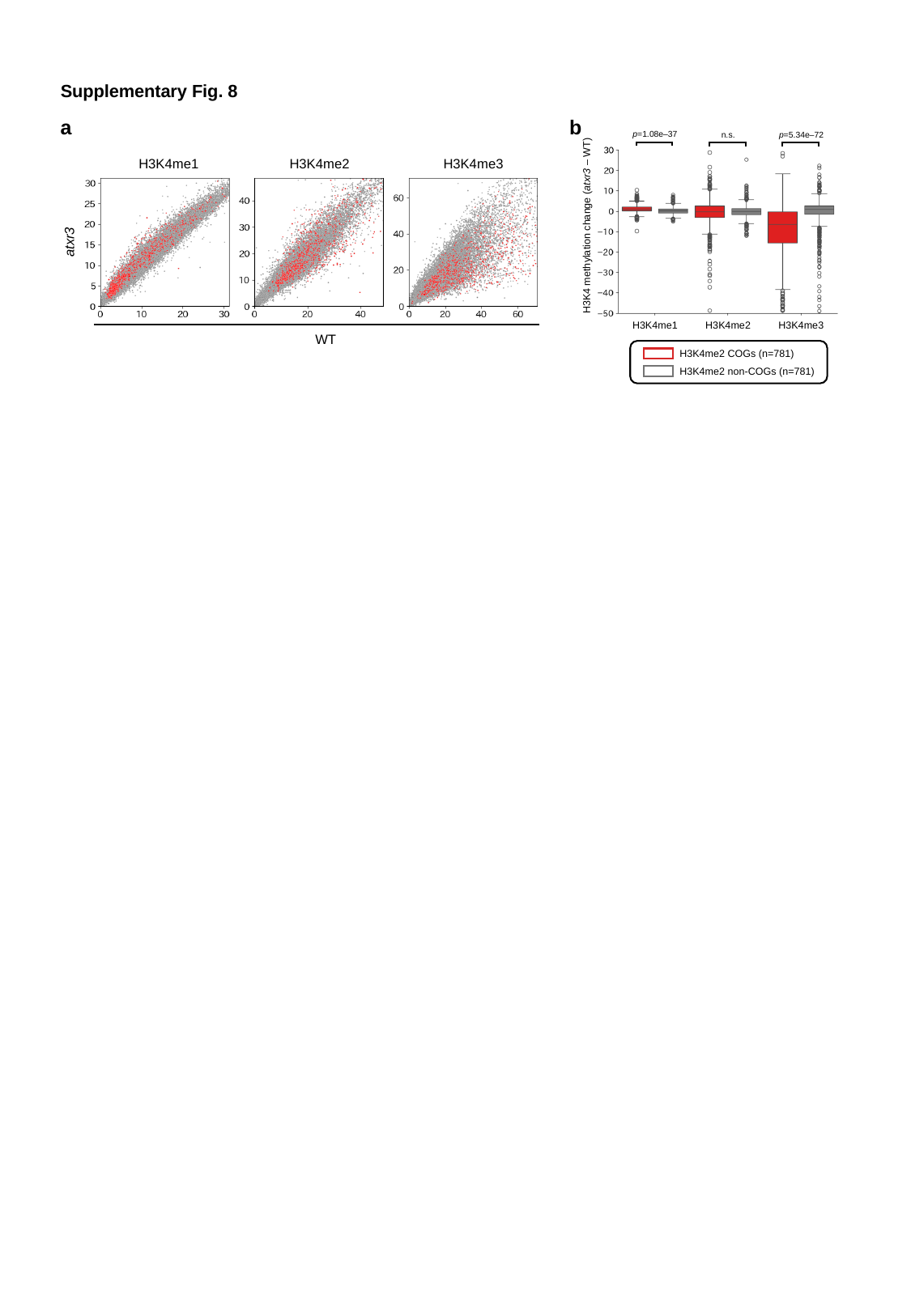

Supplementary Fig. 8
a
b
p=1.08e–37
n.s.
p=5.34e–72
H3K4 methylation change (atxr3 – WT)
H3K4me1
H3K4me2
H3K4me3
H3K4me1
H3K4me2
H3K4me3
atxr3
WT
H3K4me2 COGs (n=781)
H3K4me2 non-COGs (n=781)

### Slide 9
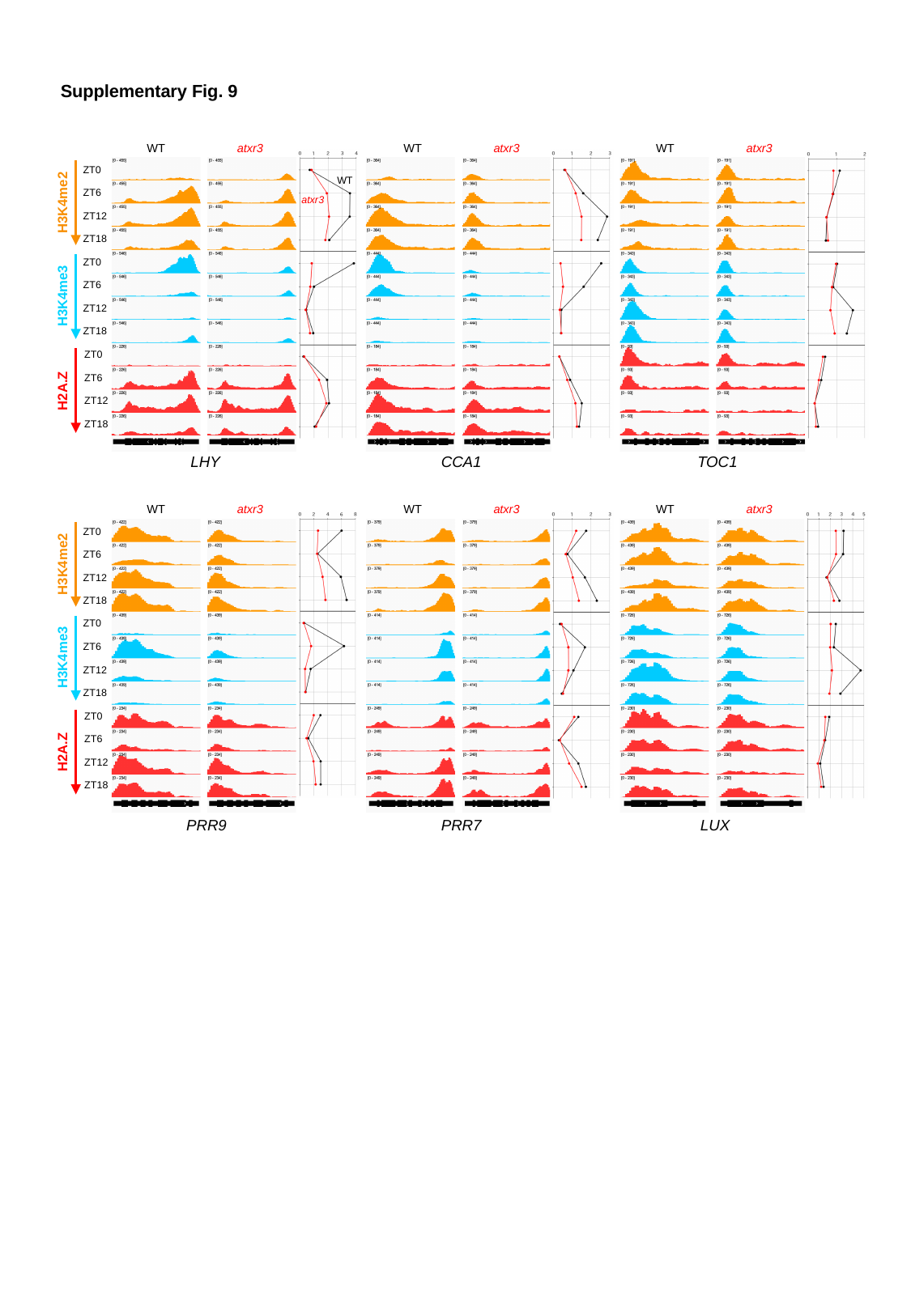

Supplementary Fig. 9
WT
atxr3
WT
atxr3
WT
atxr3
ZT0
ZT6
ZT12
ZT18
ZT0
ZT6
ZT12
ZT18
ZT0
ZT6
ZT12
ZT18
H3K4me2
H3K4me3
H2A.Z
WT
atxr3
CCA1
TOC1
LHY
WT
atxr3
WT
atxr3
WT
atxr3
ZT0
ZT6
ZT12
ZT18
ZT0
ZT6
ZT12
ZT18
ZT0
ZT6
ZT12
ZT18
H3K4me2
H3K4me3
H2A.Z
PRR7
LUX
PRR9

### Slide 10
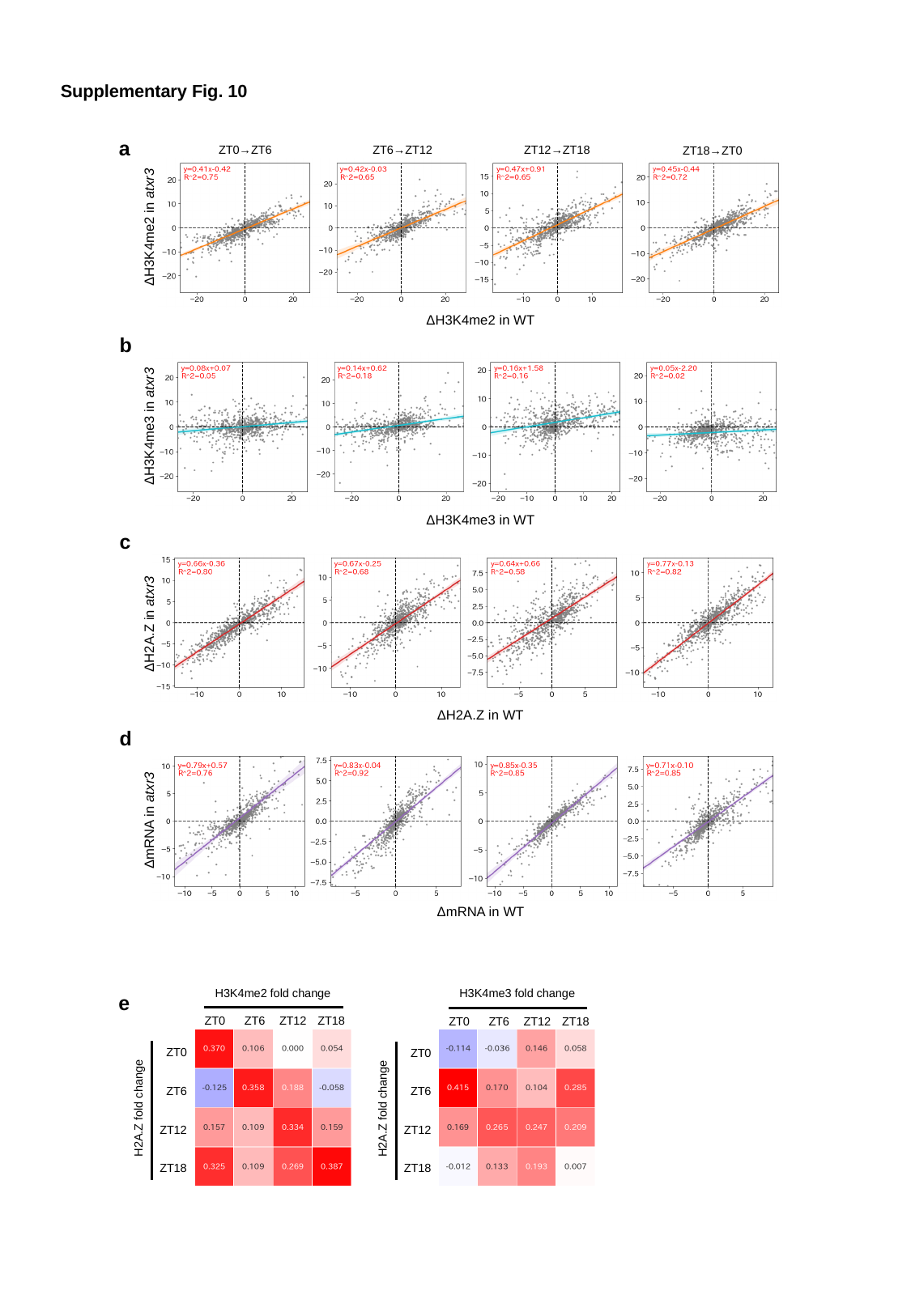

Supplementary Fig. 10
a
ZT12→ZT18
ZT0→ZT6
ZT6→ZT12
ZT18→ZT0
ΔH3K4me2 in atxr3
ΔH3K4me2 in WT
b
ΔH3K4me3 in atxr3
ΔH3K4me3 in WT
c
ΔH2A.Z in atxr3
ΔH2A.Z in WT
d
ΔmRNA in atxr3
ΔmRNA in WT
H3K4me2 fold change
ZT0
ZT6
ZT12
ZT18
ZT0
ZT6
H2A.Z fold change
ZT12
ZT18
H3K4me3 fold change
ZT0
ZT6
ZT12
ZT18
ZT0
ZT6
H2A.Z fold change
ZT12
ZT18
e

### Slide 11
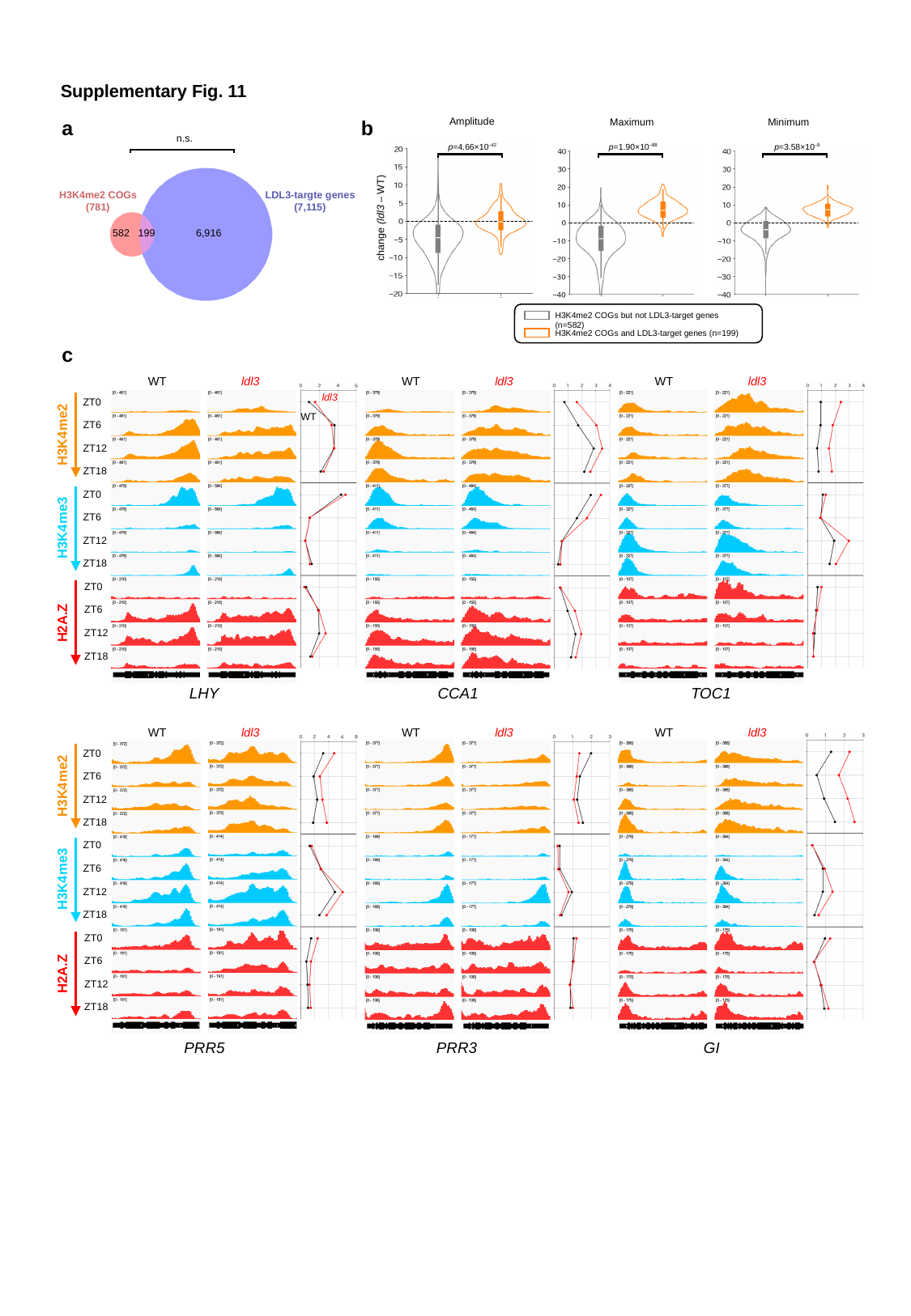

Supplementary Fig. 11
Amplitude
a
b
Minimum
Maximum
n.s.
H3K4me2 COGs
(781)
LDL3-targte genes
(7,115)
582
199
6,916
p=4.66×10^–42^
p=1.90×10^–88^
p=3.58×10^–9^
change (ldl3 – WT)
H3K4me2 COGs but not LDL3-target genes (n=582)
H3K4me2 COGs and LDL3-target genes (n=199)
c
WT
ldl3
WT
ldl3
WT
ldl3
ldl3
ZT0
ZT6
ZT12
ZT18
ZT0
ZT6
ZT12
ZT18
ZT0
ZT6
ZT12
ZT18
H3K4me2
H3K4me3
H2A.Z
WT
CCA1
TOC1
LHY
WT
ldl3
WT
ldl3
WT
ldl3
ZT0
ZT6
ZT12
ZT18
ZT0
ZT6
ZT12
ZT18
ZT0
ZT6
ZT12
ZT18
H3K4me2
H3K4me3
H2A.Z
PRR3
GI
PRR5
